# Metabolic and ribosomal dysregulation drive developmental arrest of *Schistosoma japonicum* in non-permissive rats

**DOI:** 10.1101/2025.05.20.655015

**Authors:** Yuepeng Wang, Enlu Tang, Shaoyun Cheng, Xu Chen, Wenjun Cheng, Yang Hong, Wei Hu, Jipeng Wang

## Abstract

Parasitic flatworms of the genus *Schistosoma* infect a wide range of definitive hosts, which are categorized as permissive or non-permissive based on their capacity to support parasite development. Unlike permissive hosts such as mice, *Schistosoma japonicum* undergoes pronounced developmental arrest in non-permissive hosts like rats; however, the molecular mechanisms underlying this phenomenon remain largely unclear. In this study, we conducted a comprehensive morphological and transcriptomic comparison of *S. japonicum* derived from mouse and rat hosts at multiple time points post-infection, identifying the period between 12 and 24 days post-infection (dpi) as a critical window for developmental arrest in parasites from rats. Utilizing single-cell RNA sequencing, we constructed a high-resolution cellular atlas of *S. japonicum* at 14 and 24 dpi from both host species, revealing host-dependent alterations in cell composition and transcriptional programs during parasite development. Functional analyses further demonstrated that impairments in mitochondrial function and anaerobic glycolysis contribute to the developmental arrest observed in parasites from rats. Additionally, downregulation of ribosomal genes was associated with reduced protein synthesis and impaired cell proliferation. Together, these findings suggest that disruptions in energy metabolism and ribosomal function are key drivers of developmental arrest in non-permissive hosts. This study provides novel, parasite-centered insights into host permissiveness and identifies potential molecular targets for schistosomiasis control strategies.

## Introduction

Schistosomiasis is a neglected tropical disease affecting both humans and animals, caused by parasitic flatworms of the genus *Schistosoma*. It is estimated that approximately 250 million people across 78 countries and regions are infected, with around 800 million individuals at risk and in need of preventive treatment^[1, 2]^. The schistosome life cycle involves two distinct hosts: an intermediate snail host, where the parasites undergo asexual reproduction, and a definitive mammalian host, in which they develop, undergo sexual differentiation, and produce eggs through sexual reproduction. The eggs laid by mature females are the primary driver of disease pathology in both humans and other mammals. These eggs secrete antigens at their sites of deposition—typically the liver, intestines, or bladder—eliciting host immune responses that lead to granuloma formation and subsequent fibrosis^[3-5]^. Conversely, eggs that are successfully excreted from the host hatch in pond water, releasing miracidia that infect snails and perpetuate the cycle of transmission.

To achieve sexual maturity and initiate egg production within the definitive mammalian host, schistosomes must undergo a complex migratory journey and developmental progression^[6]^. Among the three major human-infecting species, *Schistosoma japonicum* exhibits the most rapid growth rate. Following penetration of the host’s skin, *S. japonicum* cercariae transform into schistosomula within minutes and reach the subcutaneous vasculature as early as two hours post-infection^[7]^. By two hours post-infection, over 50% of schistosomula have traversed the dermis^[8]^. Within 2-3 days post-infection (dpi), the schistosomula reach the lungs, where they elongate to pass through narrow capillaries^[9]^. Subsequently, they migrate from the lungs to the portal venous system, transitioning into the liver-stage schistosomula^[9, 10]^. Notably, between 10-15 dpi, schistosomes complete the formation of a syncytial tegument, a dynamic structure essential for immune evasion, signal transduction, nutrient absorption, and continuous renewal^[11, 12]^. During this stage, the parasites attach to the vessel walls via their ventral suckers and undergo rapid growth and morphological differentiation^[13]^. Between 10-14 dpi, the worms begin to exhibit sexual dimorphism and initiate pairing behavior^[10]^. Upon pairing, the female reproductive organs, including the ovary and vitellaria, progressively develop under the influence of the male-derived pheromone BATT^[14, 15]^. Concurrently, the male reproductive system also undergoes maturation. By 24-26 dpi, the reproductive systems of both sexes are fully developed, and the adult worm pairs migrate to the mesenteric and superior rectal veins, where they commence egg deposition^[16]^.

Another significant physiological transition that occurs in schistosomes following penetration of the definitive host is the shift in respiratory metabolism. During the free-living cercarial stage, schistosomes primarily utilize aerobic respiration, metabolizing glycogen as their main energy source^[17]^. However, within hours of invading the mammalian host, a rapid metabolic shift to anaerobic respiration is initiated^[18]^. This transition enables the parasites to absorb host-derived glucose through tegument-localized *Schistosoma* glucose transporter proteins (SGTPs)^[19, 20]^. In *S. japonicum*, 23 highly homologous genes encoding glycolytic enzymes have been identified, supporting a complete anaerobic glycolytic pathway that produces lactate as the predominant end product^[18, 21]^. Interestingly, adult worms also express a subset of genes associated with aerobic metabolism, suggesting that schistosomes retain the capacity for both aerobic and anaerobic energy production depending on physiological demands^[22]^. During energetically demanding processes—such as muscular activity or tegumental repair—stored glycogen is intermittently mobilized to meet energy requirements^[18]^. This successful metabolic reprogramming, encompassing both the switch in respiratory pathways and the efficient utilization of host-derived nutrients, is considered essential for schistosome survival and adaptation within the mammalian host environment.

Schistosomes exhibit a broad host range, naturally infecting a variety of rodents, domestic animals, and livestock in endemic regions^[23]^. Notably, *S. japonicum* has been reported to infect over 40 mammalian species. However, the susceptibility of different hosts and the developmental progression of the parasite vary considerably^[24]^, allowing classification into permissive and non-permissive hosts. In non-permissive hosts, schistosome infections are either not successfully established, or the parasites fail to progress beyond the early schistosomulum stage, resulting in impaired development into adult worms, reduced egg production, or the failure of eggs to hatch into miracidia. In permissive hosts such as laboratory mice (*Mus musculus*), *S. japonicum* reaches full sexual maturity and egg-laying by approximately 35 dpi, with prominent hepatic granulomas and liver fibrosis typically observed by 42 dpi. In contrast, in non-permissive hosts such as the rat (*Rattus norvegicus*), parasites exhibit significant developmental arrest^[25, 26]^. Studies have shown that elevated expression of the *inducible nitric oxide synthase* (*iNOS*) gene in wild-type Sprague-Dawley (SD) rats impairs worm growth and reproductive development by increasing nitric oxide (NO) production. Elevated NO levels induce endoplasmic reticulum stress in schistosomes, trigger calcium efflux, and lead to mitochondrial dysfunction^[27, 28]^. In addition, NO also enhances T cell-mediated immune responses; notably, immunodeficient rats fail to restrict worm development^[29, 30]^. Furthermore, heat-sensitive components of the innate immune system, including complement component C2 and factor B of the alternative complement pathway, have also been implicated in reducing schistosome survival^[31]^. Thus, multiple factors contribute to the developmental arrest of schistosomes in non-permissive hosts such as rats.

Previous studies have predominantly focused on host-related factors—such as immune responses and biochemical conditions—in determining the permissiveness of schistosome infections. However, the parasite’s own developmental response to non-permissive host environments remains poorly understood. In this study, we systematically evaluated the developmental status of *S. japonicum* in both permissive (mouse) and non-permissive (rat) hosts at multiple time points. Morphological assessments and transcriptomic analyses revealed that the divergence in parasite development between the two hosts becomes evident between 12 and 24 dpi. Utilizing single-cell RNA sequencing, we generated a dynamic cellular atlas of *S. japonicum* in mice and rats at 14 and 24 dpi, which revealed distinct cellular impairments associated with developmental arrest in the non-permissive rat host. Further functional analyses identified disrupted energy metabolism and impaired ribosomal function as major contributors to the arrested development of *S. japonicum* in rats. These findings provide new insights into the molecular mechanisms underlying schistosomes’ host-specific developmental outcomes and suggest potential targets for therapeutic intervention and schistosomiasis control.

## Results

### Impaired development of *Schistosoma japonicum* in non-permissive rat hosts

To assess the developmental status of *Schistosoma japonicum* in the non-permissive host (Sprague-Dawley [SD] rats) compared to the permissive host (C57BL/6 [C57] mice), we infected both hosts with cercariae and recovered parasites at 12, 24, and 42 dpi **(Fig 1A)**. In mice, schistosomes recovered at 12 dpi were at the early liver stage, characterized by small size with an average worm length of approximately 500 μm. By 24 dpi, the parasites had grown substantially, reaching an average length of 6000 μm, and exhibited clear sexual dimorphism with the formation of male-female worm pairs. At 42 dpi, the worms achieved full maturity, with lengths averaging 10000-12000 μm **(Fig 1B-D)**. In contrast, schistosomes recovered from rats displayed comparable sizes to those from mice at 12 dpi. However, although male and female worms continued to develop by 24 dpi, their growth was markedly impaired, with males and females exhibiting approximately 40% and 50% reductions in length, respectively, compared to worms from mice. By 42 dpi, the size of worms from rats remained similar to that observed at 24 dpi, suggesting a developmental arrest around 24 dpi **(Fig 1B-D)**.

**Fig 1.**
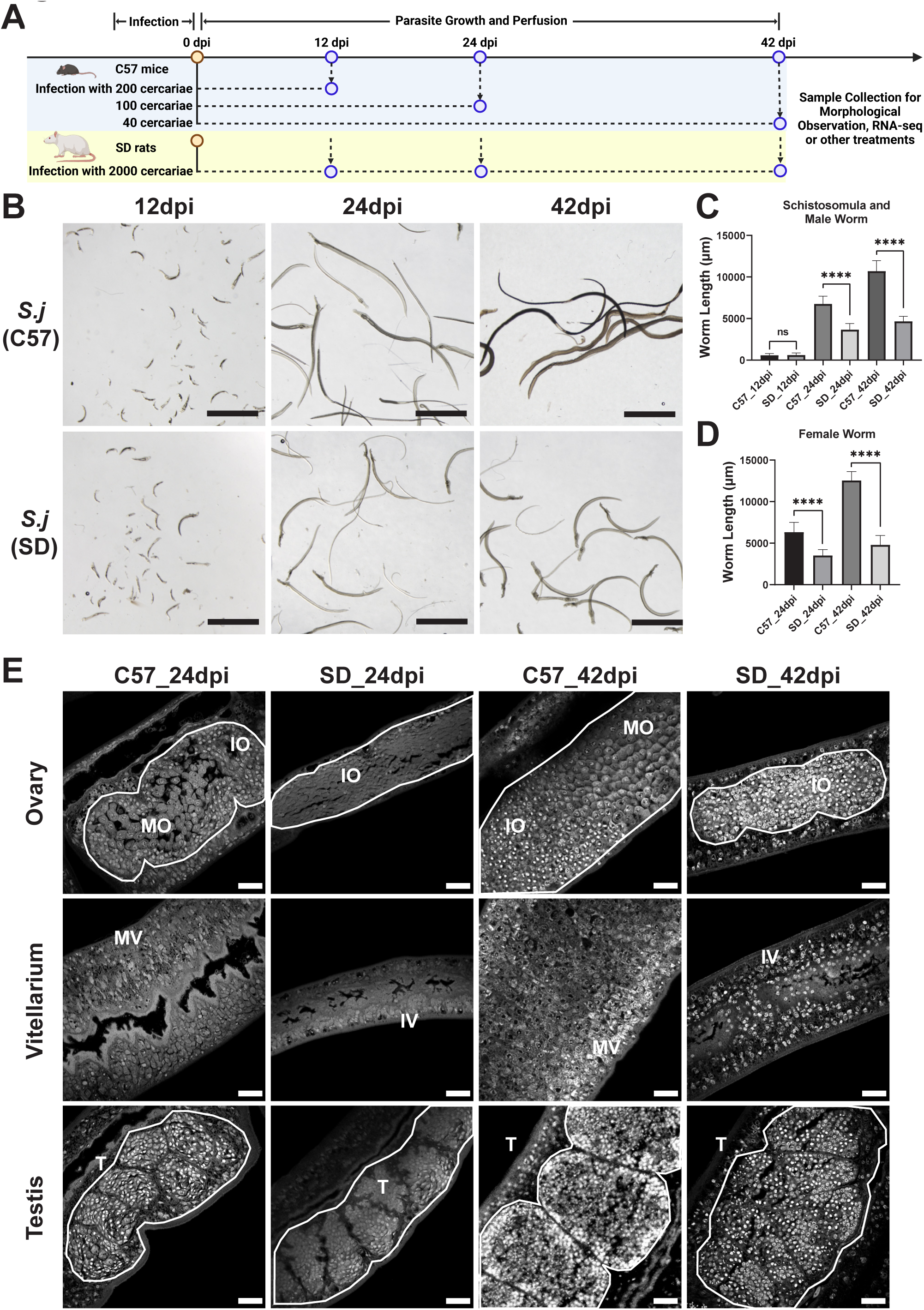
Comparative analysis of growth and reproductive development of *S. japonicum* in mice and rats. **A**, Schematic diagram illustrating the infection protocol and the timeline of parasite collection. **B**, Morphology of *S. japonicum* recovered from mice and rats at 12, 24, and 42 dpi. Scale bars, 2000 μm. **C**, Length of schistosomula and male worms from different hosts at different stages. n ≥ 12 parasites from more than three infected mice or rats. **D**, Quantification of female worm lengths from different hosts at various stages. n ≥ 14 parasites from more than three infected mice or rats. **E**, Confocal microscopy images showing the reproductive organs of male and female *S. japonicum* from mice and rats at 24 and 42 dpi. Scale bars, 25 μm. Abbreviations: MO, mature oocytes; IO, immature oocytes; MV, mature vitellocytes; IV, immature vitellocytes; T, testes; *S.j*, *S. japonicum*; C57, C57 mice; SD, SD rats. Statistical analyses were performed using independent-sample *t*-tests. ****, *p* ≤ 0.0001; ***, *p* ≤ 0.001; **, *p* ≤ 0.01; *, *p* ≤ 0.05; ns, not significant (*p* > 0.05).

We further examined the reproductive organ development of parasites recovered from mice and rats at 24 and 42 dpi. As illustrated in **Figure 1E**, parasites from mice displayed well-developed reproductive organs at 24 dpi, including mature oocytes within the ovary, mature vitellocytes in the vitellaria, and abundant spermatozoa in the testes. Conversely, the reproductive systems of parasites from rats remained immature: female worms possessed ovaries predominantly composed of immature oocytes, and mature vitellocytes were absent in the vitellaria. Male worms from rats exhibited testes with irregular, indented edges, in contrast to the mice-derived worms, whose testes comprised 7-8 well-organized, ovoid lobes with smooth margins. At 42 dpi, mouse-derived schistosomes exhibited further maturation of reproductive structures, characterized by enlarged organs and a high proportion of mature gametes. In contrast, schistosomes from rats remained developmentally arrested, with reproductive systems resembling those observed at 24 dpi.

Given the critical role of the syncytial tegument in host-parasite interactions and its dynamic morphological and biological remodeling within definitive mammalian hosts, we compared the tegumental structures of parasites from mice and rats. Scanning electron microscopy (SEM) analysis revealed that at 24 dpi, the ventral tegument of schistosomes from mice exhibited a loosely porous, interwoven structure with regularly distributed sensory papillae **(Fig 2A)**. In contrast, worms from rats displayed a mildly swollen, denser tegument with smaller sensory papillae. By 42 dpi, schistosomes from mice developed a well-defined three-dimensional interwoven tegument with enlarged sensory papillae, whereas the tegument of parasites from rats remained largely unchanged from that observed at 24 dpi **(Fig 2A)**. Transmission electron microscopy (TEM) was employed to further investigate ultrastructural differences. At 24 dpi, worms from mice possessed a well-developed syncytial layer and two robust, evenly distributed underlying muscle layers **(Fig 2B)**. Tegumental cell bodies, characterized by high electron density, were situated deeper within the tissue and connected to the syncytium via cytoplasmic bridges. In contrast, worms from rats exhibited slightly thinner syncytial and muscle layers. By 42 dpi, the syncytial and muscle layers of parasites from mice had thickened significantly, whereas those from rats remained thin **(Fig 2B)**. Notably, no substantial surface damage or host immune cell adhesion was observed in parasites from either host.

**Fig 2.**
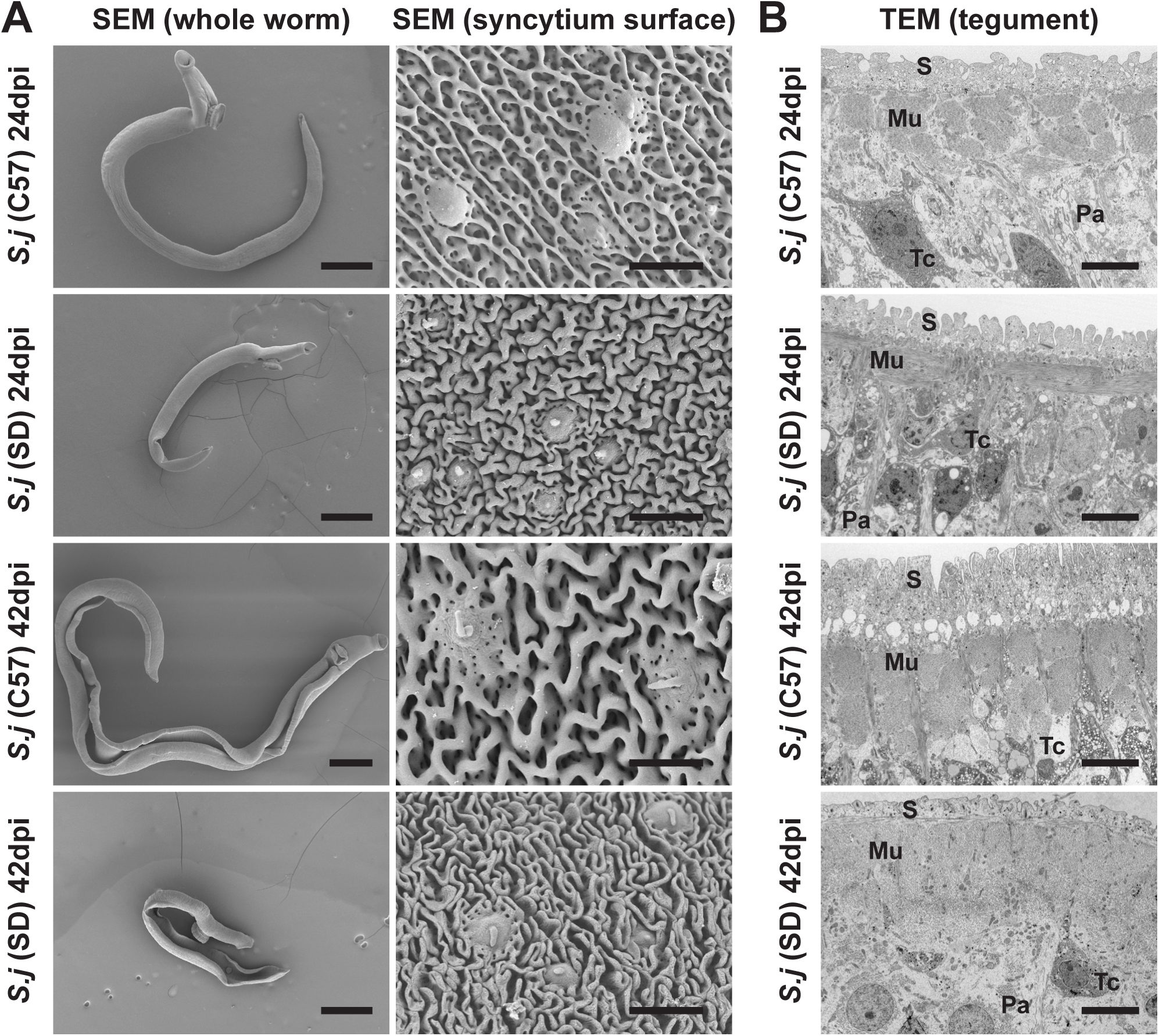
Ultrastructure of the tegument of *S. japonicum* from mice and rats. **A**, SEM images showing the overall morphology (left) and surface structure of the male parasites from mice and rats at different stages. Scale bars: 500 μm for the whole worm images, 5 μm for tegument surface images. Representative images from 3 parasites. **B**, TEM images showing the tegument and underlying tissues. Scale bars, 5 μm. Abbreviations: *S.j*, *S. japonicum*; S, syncytium; Mu, muscle; Tc, tegumental cell body; Pa, parenchyma.

Collectively, these findings suggest that multiple organ systems of *S. japonicum* are adversely affected in the non-permissive rat host, leading to impaired growth and reproductive development.

### Host-dependent transcriptomic divergence of *S. japonicum* begins between 12-24 dpi

To investigate the molecular basis of developmental arrest in *S. japonicum* in rats, we performed RNA-seq to profile gene expression patterns in parasites from mice and rats at 12, 24, and 42 dpi **(Fig 1A)**. Principal component analysis (PCA) showed that samples from mice clustered distinctly by time point and sex. As parasites developed, transcriptional differences between time points and between male and female worms became more pronounced **(Fig 3A)**. In contrast, while rat-derived worms at 12 dpi clustered separately from later stages, worms at 24 and 42 dpi clustered closely together within each sex **(Fig 3B)**. Heatmap analysis of gene expression values revealed that the transcriptional profiles of worms at 12 dpi were similar between hosts, but host-specific differences became increasingly evident from 24 dpi onward **(Fig 3C, Fig S1)**.

**Fig 3.**
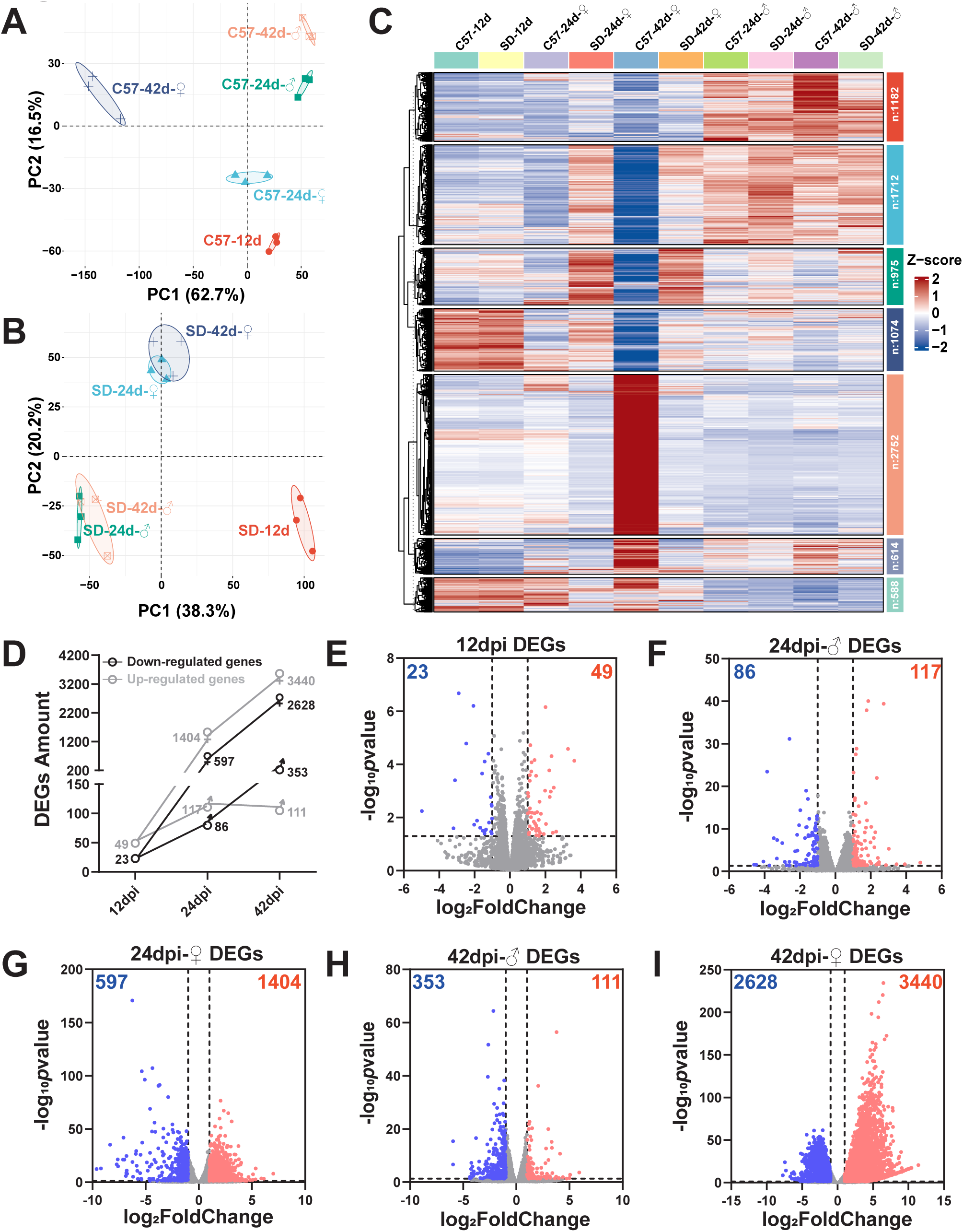
Transcriptional profile of *S. japonicum* at differential stages from mice and rates. **A-B**, PCA of parasite transcriptomes from C57 mice (**A**) and SD rats (**B**) at 12, 24, and 42dpi. **C**, *k*-means clustering of gene expression profiles for each sample group. The average gene expression across three biological replicates per group was used for clustering. **D**, Number of significantly DEGs between rat- and mouse-derived worms at 12, 24, and 42 dpi (*p* ≤ 0.05, | Log_2_FoldChange | ≥ 1). **E-I**, Volcano plots showing DEGs between worms from rats and mice at 12, 24, and 42 dpi. Abbreviations: C57, worms from C57 mice; SD, worms from SD rats.

We next assessed differential gene expression between the two hosts at each time point. At 12 dpi, only 72 genes were significantly differentially expressed (*p* < 0.05, | Log_2_FoldChange | > 1), with 49 upregulated and 23 downregulated in rat-derived worms **(Fig 3D, E, Table S1)**. The number of differentially expressed genes (DEGs) increased with parasite development, with female worms exhibiting both a higher number of DEGs and greater fold changes compared to males **(Fig 3D-I, Table S1)**, likely due to the substantial contribution of sexual organs to the body size in mature females. Consistent with our previous morphological observations, these transcriptomic analyses demonstrate that worms at 12 dpi share a similar developmental status in both hosts. However, from 12-24 dpi, parasites begin to exhibit strong host-dependent transcriptional divergence, suggesting that host-induced growth arrest initiates during this period.

To identify the cell types affected in the non-permissive host, we mapped the significantly downregulated genes from *S. japonicum* worms isolated from rats onto our previously constructed single-cell atlas of adult *S. japonicum*. At 12 dpi, the downregulated genes were predominantly enriched in proliferative cell populations, including somatic stem cells, tegument progenitor cells, and neuronal stem cells **(Fig S2)**. By 24 dpi, the downregulated genes were primarily associated with differentiated cell types. In male worms, these genes were mainly localized to the parenchyma, muscle, and tegument progeny 1/2 cell clusters, whereas in female worms, they were highly enriched in reproductive organ cells **(Fig S2)**. Collectively, these findings suggest that the developmental arrest of *S. japonicum* in rats may result from impairments in proliferative cells, subsequently leading to defects across multiple tissues during later stages of development.

### Single-cell transcriptomics reveals cellular defects in arrested *S. japonicum* development in rats

To elucidate the cellular basis of developmental arrest in *S. japonicum* within rats, we employed 10× Genomics single-cell RNA sequencing (scRNA-seq) to profile cell population dynamics during the critical period of 12-24 dpi. Given that worms exhibited no sexual dimorphism at 12 dpi, we selected 14 dpi, a stage at which male and female parasites could be distinguished, for analysis. Additionally, considering that female parasite development is strongly influenced by pairing with male worms, we focused on male worms for scRNA-seq, collecting samples from both mice and rats at 14 and 24 dpi. Using marker genes previously identified in *S. mansoni* as reference **(Table S2)**^[32]^, we classified 60 distinct cell clusters of *S. japonicum* into 17 cell types **(Fig 4A, Fig S3A, B)**

**Fig 4.**
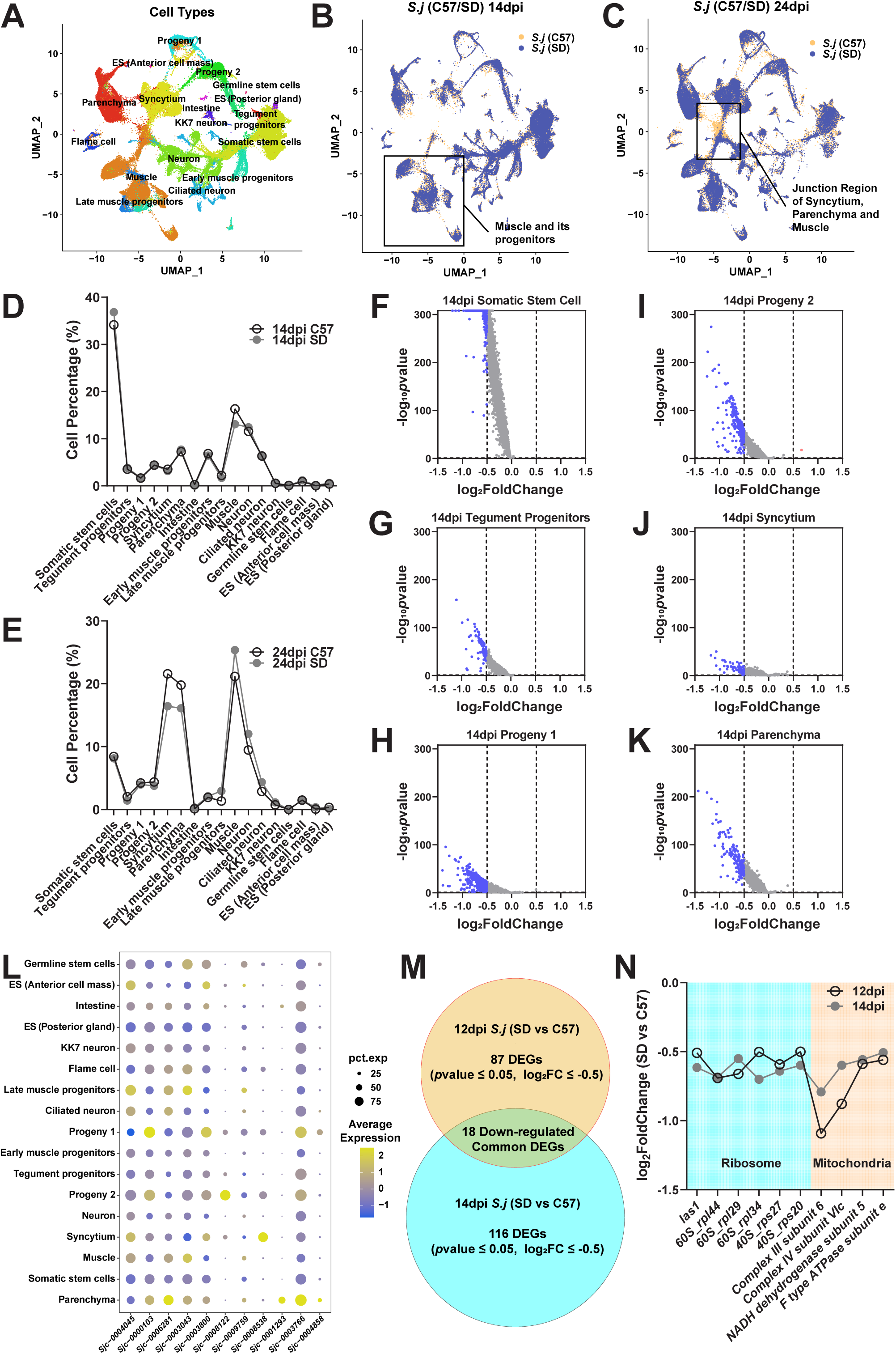
Single-cell transcriptome atlas of *S. japonicum* at differential stages in mice and rates. **A**, UMAP dimensionality reduction and clustering of integrated male parasite samples at 14 and 24 dpi from both hosts. **B-C**, Overlaid UMAP plots of male worm samples from mice and rats at 14 dpi (**B**) and 24 dpi (**C**), with notable cell types highlighted by boxed regions. **D-E**, Quantitative comparison of cell type proportions in male worms from mice and rats at 14 dpi (**D**) and 24 dpi (**E**). **F-K**, Volcano plots illustrating differentially expressed genes between male worms from mice and rats at 14 dpi across somatic stem cells, tegumental lineages (tegument progenitors, progeny 1, progeny 2, and syncytium), and parenchyma cells. **L**, Localization of major downregulated genes at the single-cell level in *S. japonicum* from rats at 14 dpi. Bubble size represents the percentage of cells expressing the corresponding gene, and bubble color indicates average expression level. **M**, Venn diagram depicting the overlap of significantly downregulated genes (*p* ≤ 0.05, | Log_2_FoldChange | ≥ 0.5) identified by bulk RNA-seq and scRNA-seq in schistosomula from rats. **N**, Expression profiles of commonly downregulated ribosomal and mitochondrial genes in worms from rats compared with those from mice. Hollow black dots represent bulk RNA-seq data; solid gray dots represent scRNA-seq data. Abbreviations: *S.j*, *S. japonicum*; ES, esophageal glands.

Comparison of UMAP plots revealed significant differences in the coverage areas and cell densities between groups, particularly in muscle cells at 14 dpi and in the junction region comprising syncytium, muscle, and parenchyma cells at 24 dpi **(Fig 4B, C, Fig S4A-D)**. Quantitative analysis further confirmed these observations: at 14 dpi, muscle cells constituted approximately 16.31% of total cells in worms from mice, compared to 13.09% in those from rats **(Fig 4D)**. At 24 dpi, the proportion of syncytium cells in worms from rats decreased from 21.60% to 16.44%, while parenchyma cells decreased from 19.79% to 16.11%, relative to worms from mice **(Fig 4E)**.

Beyond changes in cell composition, we next examined gene expression alterations within individual cell types. At 14 dpi, a substantial downregulation of gene expression was observed across multiple *S. japonicum* cell types in worms from rats. Germline stem cells exhibited the highest number of downregulated genes, followed by tegument progeny 1/2, somatic stem cells, esophageal glands, and parenchyma cells, with somatic stem cells showing the greatest statistical confidence for downregulation **(Fig S5; Fig 4F-K)**. In 24 dpi male worms from rats, downregulated genes were most prominently associated with flame cells and tegument progeny 2, whereas upregulated genes were primarily enriched in the esophageal glands, late muscle progenitors, and tegument progeny 1 **(Fig S5)**. The early downregulation of gene expression in stem and progenitor cell populations at 14 dpi is likely to have directly contributed to the impaired development of multiple cell lineages at later stages.

In cell types that constitute the predominant cellular populations within the parasite—such as somatic stem cells, the tegument lineage, and parenchymal cells **(Fig 4D)** —DEGs identified in schistosomula derived from rats exhibited substantial overlap. Notably, these included *S. japonicum L-lactate dehydrogenase A chain* (*SjL-ldh*, *Sjc_0004045*), *universal stress protein in the QAH/OAS sulfhydrylase 3’ region* (*Sjc_0006281*), *ubiquinol cytochrome C oxidoreductase subunit 6* (*Sjc_0003043*), and *actin-related protein 2/3 complex subunit 3-B* (*Arp2/3 subunit 3-B*, *Sjc_0003800*) **(Table S3, 4)**. These genes tended to exhibit high expression levels in parenchyma, progeny 2, muscle, and late muscle progenitor cells **(Fig 4L)**, suggesting that repression of gene expression in these specific cell types may contribute to the developmental arrest observed in the parasite.

To identify key genes underlying the developmental arrest of schistosomes in rats, we performed an overlapping analysis between the 116 significantly downregulated genes identified by scRNA-seq of 14 dpi male parasites and the 87 significantly downregulated genes identified by bulk RNA-seq of 12 dpi worms. This analysis revealed 18 genes that were consistently downregulated in both datasets **(Fig 4M; Table S5)**. Among these, four genes encode mitochondrial proteins associated with the electron transport chain (ETC), and six genes are involved in ribosomal function **(Fig 4N; Table S5)**. The expression of these genes was reduced by more than 1.4-fold, suggesting that in schistosomula from rats, a subset of mitochondrial and ribosomal genes was consistently downregulated and comprised a substantial portion of the stable, downregulated gene set **(Fig 4N)**.

### Energy metabolism impairment limits *S. japonicum* development in rats

We further evaluated alterations in mitochondrial gene expression in 14 dpi worms from the two different hosts. Among the 12 mitochondrial DNA-encoded genes, significant downregulation of *NADH dehydrogenase subunits 1–6* (*nd1–6*) and *cytochrome c oxidase subunit 1* (*cox1*) was observed in parasites from rats across all cell populations, with the exception of intestinal cells **(Fig S6)**. These downregulated mitochondrial genes were predominantly expressed in high energy-demanding tissues, including muscle and its progenitors (early and late muscle progenitors) in schistosomula **(Fig 5A)**. The suppression of mitochondrial ETC genes is likely to impair the energy supply derived from aerobic respiration, thereby compromising parasite viability in rats. To validate this hypothesis, 14 and 24 dpi worms were treated *in vitro* for 48 hours with the mitochondrial membrane potential uncouplers oligomycin and FCCP. Treatment with either uncoupler induced mitochondrial gene downregulation similar to that observed in worms from rats **(Fig S7)** and led to a marked reduction in worm viability, characterized by an inability to adhere to the culture dish via oral and ventral suckers, decreased motility, body contraction, and abnormal twisting movements **(Fig 5B)**.

**Fig 5.**
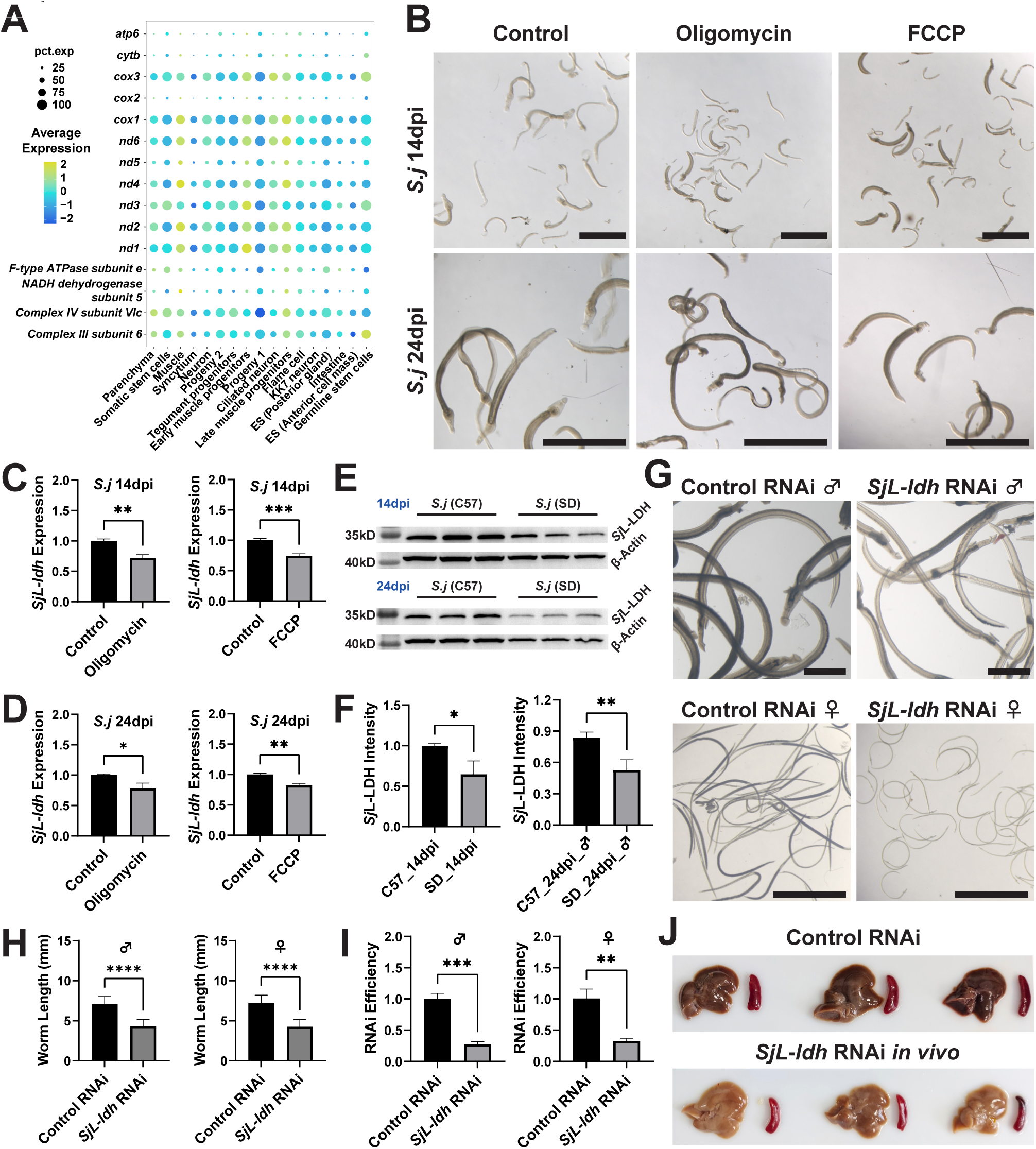
Inhibition of mitochondrial ETC and anaerobic glycolysis drives *S. japonicum* developmental arrest. **A**, Cell type-specific expression of mitochondrial genes in 14 dpi worms. Bubble size indicates the percentage of cells expressing each gene; color represents average expression level. **B**, Brightfield images of 14 and 24 dpi worms after 48 h *in vitro* treatment with oligomycin or FCCP. Scale bars, 2000 μm. **C-D**, *SjL-ldh* expression levels after oligomycin or FCCP treatment on worms at 14 dpi and 24 dpi. **E**, Western blot analysis of *Sj*L-LDH and β-actin in worms from mice and rats at 14 and 24 dpi. **F**, Quantification of Western blot showing relative *Sj*L-LDH expression between hosts. **G**, Brightfield images of worms from control and *SjL-ldh in vivo* RNAi groups in mice. Scale bars, 1000 μm. **H**, Worm length comparison between control and *SjL-ldh in vivo* RNAi groups. **I**, RNAi efficiency of *SjL-ldh* knockdown in male and female worms. **J**, Gross morphology of livers and spleens from control and *SjL-ldh in vivo* RNAi mice. Abbreviations: ES, esophageal glands. Statistical analyses were performed using independent-sample *t*-tests. ****, *p* ≤ 0.0001; ***, *p* ≤ 0.001; **, *p* ≤ 0.01; *, *p* ≤ 0.05; ns, not significant (*p* > 0.05).

In the definitive host, *S. japonicum* undergoes a metabolic shift from aerobic glucose metabolism to anaerobic glycolysis, with lactate serving as the primary end product^[18, 21, 33]^. *SjL-ldh*, encoding L-lactate dehydrogenase, an enzyme that catalyzes the reversible conversion between pyruvate and lactate^[34]^, was found to be downregulated across multiple tissues of schistosomula in rats **(Fig 4L, Table S3)**. qPCR analysis following 48-hour treatment with mitochondrial uncouplers demonstrated that both FCCP and oligomycin induced an approximately 25% reduction in *SjL-ldh* expression in male parasites at 14 and 24 dpi from mice **(Fig 5C, D)**. These findings suggest that mitochondrial dysfunction may impair the anaerobic glycolytic pathway in *S. japonicum*, thereby further inhibiting parasite growth and development by limiting energy metabolism in the rat host.

According to our single-cell atlas, *SjL-ldh* was broadly expressed throughout the parasite, with particularly high expression in energy-demanding cell types such as the syncytium, tegument progenitor cells, muscle cells, and late muscle progenitor cells **(Fig S8)**. These tissues are likely to have a greater reliance on anaerobic respiration, given the high turnover of the syncytial tegument and the critical role of muscle in parasite motility^[18]^. Furthermore, scRNA-seq analysis revealed that 21 glycolysis-related genes, 26 genes associated with the tricarboxylic acid (TCA) cycle, and 49 genes involved in the ETC and oxidative phosphorylation were differentially expressed in 14 dpi schistosomula recovered from rats. Among these, *SjL-ldh* exhibited the most pronounced downregulation **(Table S6)**. Given the high sequence similarity between the *SjL-ldh* A chain and homologous lactate dehydrogenases from other model organisms, including humans and mice **(Table S7, Fig S9)**, we employed a commercially available antibody against a synthetic peptide corresponding to a region of LDH for western blot analysis to assess *Sj*L-LDH protein levels in *S. japonicum* from rats and mice at different stages. *Sj*L-LDH protein levels were approximately 35% and 37% lower in worms from rats compared to those from mice at 14 dpi and 24 dpi, respectively **(Fig 5E, F)**. The consistent reduction in *SjL-ldh* expression at both mRNA and protein levels indicates that anaerobic respiration is significantly impaired in *S. japonicum* in rats.

To investigate whether inhibition of anaerobic respiration affects the development of *S. japonicum*, we conducted RNA interference (RNAi) targeting *SjL-ldh in vivo*. Following 24 days of *SjL-ldh* dsRNA treatment in mice, both male and female worms exhibited impaired development, characterized by an approximate 40% reduction in body length, while worm burden remained unaffected **(Fig 5G, H, Fig S10)**. Quantitative PCR analysis confirmed a significant suppression of *SjL-ldh* expression by RNAi **(Fig 5I)**. Additionally, *SjL-ldh* knockdown alleviated pathological damage in the livers and spleens of treated mice **(Fig 6J)**. Further analysis of mitochondrial ETC-related DEGs revealed that silencing *SjL-ldh* did not alter the expression of these mitochondrial genes **(Fig S11)**, suggesting that the glycolytic disruption resulting from *SjL-ldh* suppression in worms from rats is likely a downstream consequence of primary mitochondrial dysfunction.

**Fig 6.**
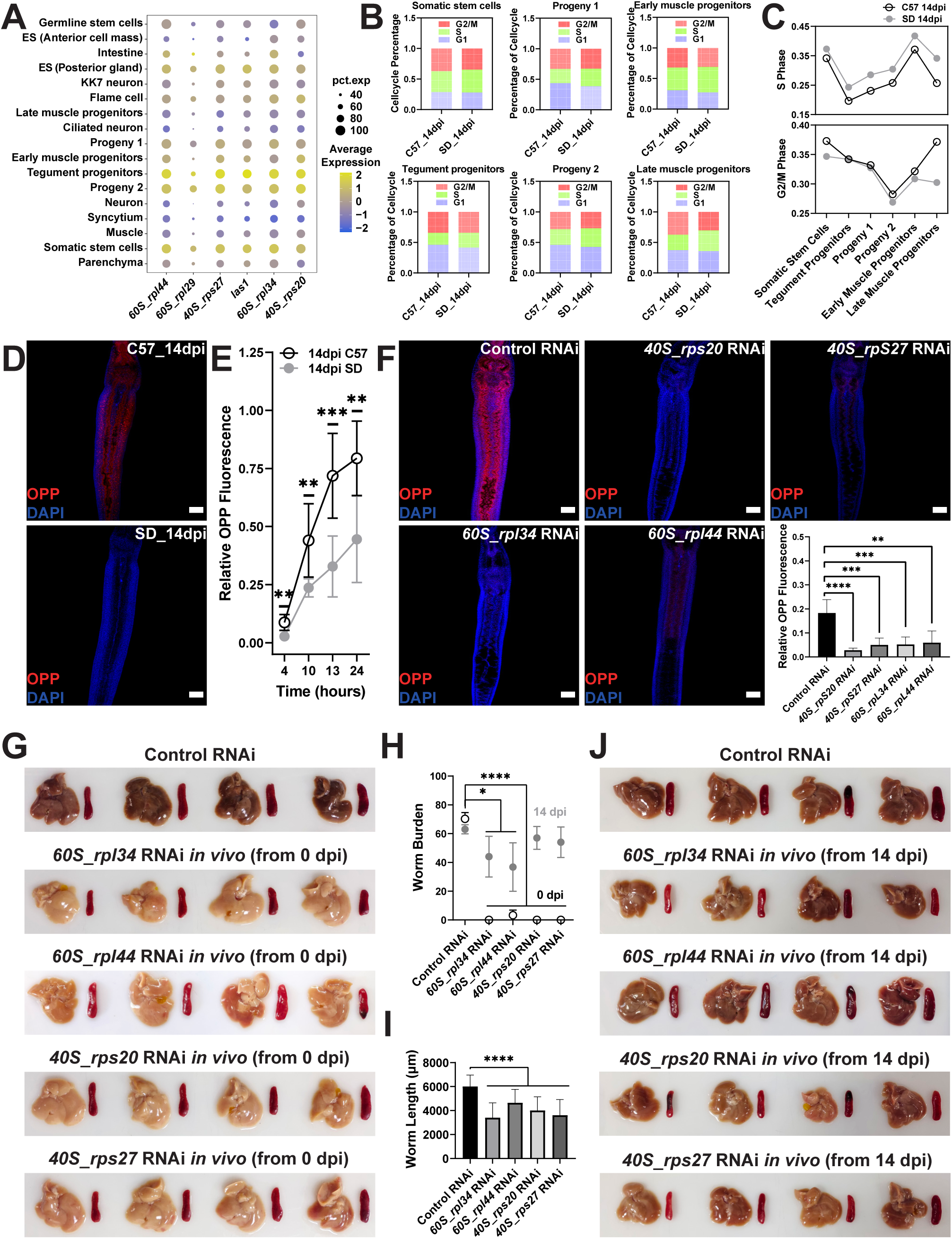
Ribosomal dysfunction reduces protein synthesis and hinders schistosome development. **A**, Cell type-specific expression of ribosomal genes in 14 dpi *S. japonicum*. Bubble size represents the percentage of cells expressing each gene; color indicates average expression level. **B-C**, Cell cycle phase prediction (**B**) and quantitative comparison (**C**) in somatic stem cells, tegument cell lineages, and parenchymal cell clusters from 14 dpi worms derived from mice and rats. **D**, OPP labeling of nascent proteins in 14 dpi worms from mice and rats after 4 h of incubation. Scale bars, 100 μm. **E**, Quantification of OPP fluorescence intensity over 24 h in 14 dpi schistosomula recovered from mice and rats. **F**, OPP labeling and quantification of nascent protein synthesis following 4 h incubation in control versus ribosomal gene RNAi-treated worms *in vitro*. Scale bars, 100 μm. **G**, Gross pathology of livers and spleens from mice treated with dsRNA against ribosomal genes starting at 0 dpi. **H**, Worm burden in mice treated with ribosomal gene dsRNA from 0 dpi and from 14 dpi. **I**, Worm length in mice treated with ribosomal gene dsRNA from 14 dpi. **J**, Gross pathology of livers and spleens from mice treated with dsRNA against ribosomal genes starting at 14 dpi. Abbreviations: C57, worms from C57 mice; SD, worms from SD rats; ES, esophageal glands. Statistical analyses were performed using independent-sample *t*-tests. ****, *p* ≤ 0.0001; ***, *p* ≤ 0.001; **, *p* ≤ 0.01; *, *p* ≤ 0.05; ns, not significant (*p* > 0.05).

### Ribosomal gene downregulation impairs protein synthesis and limits *S. japonicum* development in rats

Bulk and scRNA-seq analyses revealed that five ribosomal subunit protein genes, along with ribosome biogenesis protein *las1*, were consistently downregulated in 14 dpi schistosomula from rats **(Fig 4N)**. Meanwhile, the Gene Set Enrichment Analysis (GSEA) of samples from rat host at 12 and 24 dpi **(Table S8)**, compared with those from mice, indicated that biological functions involving ribosome activity, protein synthesis, and amino acid metabolism were significantly affected **(Fig S12)**. Given the critical role of ribosomal proteins in protein synthesis, we hypothesized that their downregulation may contribute to the developmental arrest observed in schistosomes in rats. To explore this, we first assessed the expression of these genes following *SjL-ldh* RNAi, a treatment previously shown to induce parasite developmental arrest **(Fig 5G)**. Notably, *SjL-ldh* knockdown did not affect the expression levels of ribosomal genes, suggesting that ribosomal dysfunction occurs independently of energy metabolism disruption **(Fig S13)**. Furthermore, single-cell atlas data demonstrated that these ribosomal genes are highly expressed in somatic stem cells, early muscle progenitors, tegument progeny 2 cells and tegument progenitors which are characterized by high proliferative and differentiation potential **(Fig 6A)**.

Cell proliferation is a fundamental biological process closely associated with organismal development and is critically dependent on the stability of ribosome biogenesis and function^[35, 36]^. The downregulation of ribosome-related genes may impair the proliferative capacity of schistosomula in rats. To investigate this possibility, EdU labeling was employed to track newly replicated DNA in proliferating cells within the worms. This analysis revealed no significant differences in the proportion of proliferative cells between worms from mice and rats during 12-24 dpi **(Fig S14A-C)**. Subsequently, cell cycle phase prediction was performed on proliferative cell clusters from 14 dpi scRNA-seq data using a manually curated set of S- and G2/M-phase marker genes **(Table S9)**. The results demonstrated a modest increase in the proportion of S-phase cells and a corresponding decrease in G2/M-phase cells in worms from rats relative to those from mice, with the most pronounced difference observed in late muscle progenitors **(Fig 6B, C)**. These findings suggest that proliferative cells in schistosomula from rats experience an S-G2/M phase arrest, potentially linked to ribosomal dysfunction.

To further investigate whether the down-regulation of ribosomal protein genes in worms derived from rats is associated with impaired protein synthesis, we employed O-propargyl-puromycin (OPP), an alkyne analog of puromycin, to label nascent proteins. In 14 dpi worms derived from mice, strong labeling was observed throughout the body after 4 hours, with signal intensity increasing over time and reaching a peak at 24 hours **(Fig 6D, E, Fig S15)**. In contrast, nearly no labeling was observed in 14 dpi schistosomula from rats at the 4-hour time point, although signal intensity gradually increased with extended labeling durations **(Fig 6D, E, Fig S15)**. These results suggest that protein synthesis is slower in worms derived from rats.

We next assessed the impact of ribosomal gene expression on protein synthesis by knocking down these genes individually in 14 dpi parasites derived from mice *in vitro*. Silencing each of the six target genes did not noticeably affect parasite morphology or vitality compared to the control RNAi group **(Fig S16)**. However, a significant reduction in nascent protein synthesis, as detected by OPP labeling, was observed following the knockdown of *40S-rps20*, *40S-rps27*, *60S-rpl34*, and *60S-rpl44* **(Fig 6F)**. These findings suggest that the diminished protein synthesis capacity observed in *S. japonicum* from rats is associated with the down-regulation of ribosomal gene expression.

Ribosomal dysfunction and the consequent reduction in protein synthesis may impair worm development in rats. To investigate this, we performed RNA interference (RNAi) targeting ribosome-related genes that were down-regulated in worms from rats, starting at 0 dpi in infected mice. Following several days of dsRNA treatment targeting four genes, mice exhibited markedly reduced pathological changes in both the liver and spleen compared to the control group **(Fig 6G)**. Notably, no worms were recovered from the *40S-rps20*, *40S-rps27*, and *60S-rpl34* RNAi groups, while significantly fewer worms were retrieved from the *60S-rpl44* RNAi groups, suggesting that these ribosome-associated genes are essential for schistosome survival in the host **(Fig 6H, Fig S17A)**. Given that morphological and transcriptional divergence between parasites from mice and rats emerges around 12 dpi, we subsequently initiated dsRNA treatment at 14 dpi. This later intervention led to a significant reduction in worm length and a moderate decrease in worm burden **(Fig 6H, I, Fig S17B)**, accompanied by attenuated liver and spleen pathology **(Fig 6J)**. These findings indicate that the down-regulation of ribosomal genes in worms from rats during the schistosomula stage contributes to their developmental arrest.

## Discussion

Although rats are considered non-permissive hosts for schistosomes, their hostile internal environment does not appear to affect parasite growth during the early stages of infection. Zhou et al. reported that the length of liver-stage *S. japonicum* at 14 dpi in SD rats was comparable to that in mice^[37]^. Similarly, Cioli et al. observed that the development of *S. mansoni* in rats was apparently normal during the first four weeks post-infection^[38]^. In the present study, we found that *S. japonicum* at 12 dpi exhibited morphological similarity in both mice and rats **(Fig 1B)**. This observation was further supported by bulk RNA-sequencing, which revealed only a few DEGs between parasites from the two hosts at this stage **(Fig 2D, E)**. At 24 dpi, worm length in rats was approximately half that observed in mice, accompanied by marked differences in gene expression profiles **(Fig 1B**, **Fig 2D, F, G)**. Our data indicate that the critical window for developmental arrest likely occurs between 12 and 24 dpi in rats. Unlike in rats, *S. japonicum* exhibits nearly 90% clearance within 10 dpi in another non-permissive host, *M. fortis*, accompanied by observable tegumental damage, cellular adhesion, and trogocytosis on the syncytial surface^[39, 40]^.These findings further suggest that the impact of the rat host on *S. japonicum* involves a progressive and sustained suppression beginning at the schistosomula-to-adult transition, rather than being mediated by an immediate and intense immune rejection following infection.

Consistent with previous reports, the most significantly affected tissues were the reproductive organs in both male and female worms^[41]^. Interestingly, although the testes of male worms from rats appeared to have similarly sized testicles, they exhibited a wrinkled surface and contained few secondary spermatocytes **(Fig 1E)**. In contrast, female reproductive organs in rats remained largely undeveloped at both 24 and 42 dpi **(Fig 1E)**. It is well established that female reproductive development in schistosomes relies on sustained pairing stimulation from males, which is mediated by the transfer of male pheromones through the gynecophoric canal, presumably via the ciliated neurons (sensory papillae) located on the tegument^[14, 15]^. In this study, we observed that the expression of *nrps* (*Sjc_0000814*), which encodes the synthetic enzyme responsible for male pheromone production^[14, 15]^, was significantly downregulated in male worms from rats, showing a 4.1-fold reduction at 24 dpi) and a 1.8-fold reduction at 42 dpi (**Table S1**). Moreover, significant structural differences were observed in the ventral tegument of male worms from rats compared to those from mice at 24 dpi **(Fig 2A)**. Notably, the sensory papillae of male worms in rats were smaller and malformed **(Fig 2A)**. These structural alterations may not only impair the male’s ability to sense female for the upregulation of *nrps* and subsequent pheromone synthesis, hinder the effective transmission of pheromones from male to female. Consequently, these disruptions could contribute to impaired female sexual maturation, despite the continued occurrence of male-female pairing behavior. These findings indicate that the development of multiple organs in *S. japonicum* is impaired in the non-permissive rat host, ultimately restricting overall worm growth.

Several studies have investigated the gene expression profiles of *S. japonicum* in non-permissive hosts using microarray and bulk RNA-seq approaches^[26, 42, 43]^. However, these conventional methods are limited in their ability to capture cellular heterogeneity in gene expression. In the present study, we characterized for the first time the transcriptional changes in *S. japonicum* at single-cell resolution using scRNA-seq of male worms derived from mice and rats at key developmental stages. This single-cell atlas revealed that the effects of the rat host on the parasite were not uniformly distributed across all worm cell types, but were predominantly concentrated in somatic stem cells and progenitor populations during the schistosomulum stage **(Fig 4D-K)**. The somatic stem cells, or neoblasts, exhibit substantial heterogeneity and possess the capacity to differentiate into various cell types, including those of the tegument, gut, and muscle^[44, 45]^. Notably, experimental ablation of stem cells through irradiation disrupts tissue repair and renewal, ultimately leading to parasite clearance within the host^[45, 46]^. Given that schistosome growth and development are driven by stem cell populations, disruptions in these cells have profound consequences. In this study, the sustained suppression of gene expression in proliferative cells ultimately impaired the development of tegumental lineages, parenchymal tissues, and other cell types, thereby further inhibiting parasite growth in rats.

Throughout their complex life cycle, schistosomes undergo dynamic shifts in respiratory metabolism to adapt to distinct environmental conditions^[18, 33]^. During the free-living cercarial stage, the parasite relies on stored glycogen and primarily utilizes aerobic respiration for ATP production^[17, 33]^. Upon invasion of the definitive host, *S. japonicum* rapidly switches to anaerobic metabolism, utilizing host-derived glucose to support energy-intensive processes such as reproduction and repair of the syncytial tegument^[18, 47]^. In this study, we observed a significant downregulation of mitochondrial genes, particularly those expressed in energy-demanding tissues such as muscle and its progenitor cells, leading to impaired function of the ETC involved in aerobic respiration **(Fig 4N**, **Fig 5A, Fig S6)**. Pharmacological disruption of the ETC using the uncoupling agents FCCP and oligomycin resulted in the downregulation of mitochondrial gene expression and a marked reduction in worm viability, supporting a direct link between mitochondrial gene suppression and developmental arrest **(Fig 5B, Fig S7)**. Furthermore, ETC dysfunction was associated with reduced expression of the glycolytic gene *SjL-ldh*, which was among the most significantly downregulated glycolytic enzymes across multiple cell types **(Table S3, Fig 5C, D)**. Silencing *SjL-ldh* in mice replicated the developmental arrest phenotype observed in worms from rats, without affecting mitochondrial gene expression **(Fig 5G-I, Fig S11)**. Elevated LDH expression is typically indicative of high cellular proliferation rate^[48, 49]^. Proliferative stem cells often rely on LDH-mediated conversion of pyruvate to lactate to meet energy demands, supporting both maintenance of stemness and differentiation into mature cell lineages^[50, 51]^. Accordingly, inhibition of LDH has been proposed as a strategy to target cancer stem cells^[52, 53]^. The sustained downregulation of *SjL-ldh* observed in schistosomes from rats may thus reflect impaired differentiation capacity and proliferative arrest. Previous ultrastructural analyses have reported mitochondrial stress in worms from rats, characterized by organelle swelling, inner membrane disruption, cristae disorganization, and vacuolization^[28, 41]^. Our findings suggest that such structural damage compromises ETC function, leading to reduced *SjL-ldh* expression and diminished energy metabolism via respiratory pathways, ultimately contributing to developmental arrest. Given the central role of mitochondrial components and glycolytic enzymes in schistosome development and viability, we propose these pathways as promising targets for therapeutic intervention.

Ribosome biogenesis was a fundamental metabolic process in proliferative cells, playing a pivotal role in dynamically regulating ribosome abundance in stem cells^[35, 36]^. The maintenance of ribosomal quantity and composition, alongside the preservation of translational fidelity, is tightly coordinated to support cellular growth, proliferation, differentiation, and overall organismal development^[54]^. This coordination ensures a sufficient supply of proteins for physiological functions and accommodates the heightened demand for de novo protein synthesis during the G1 and G2 phases of the cell cycle, thereby contributing to cell cycle progression^[55, 56]^. Disruptions in ribosome assembly or alterations in ribosomal protein composition are known to induce cell cycle arrest at the G1-S or G2/M transition, potentially resulting in apoptosis^[57, 58]^. Across diverse organisms, structural deficiencies and assembly defects in ribosomes markedly impair cell proliferation and compromise normal development^[59-63]^. In this study, we observed a consistent down-regulation of multiple ribosomal protein genes in schistosomula isolated from rats **(Table S5, Fig 4N)**. Cell cycle prediction based on scRNA-seq data suggested that the proliferative cells of *S. japonicum* in rats likely undergo arrest at the S-G2/M phase **(Fig 6B, C)**. Additionally, we detected a significant reduction in protein synthesis capacity in schistosomula derived from rats or following RNAi **(Fig 6D-F, Fig S15)**. Given the critical role of ribosome function in cellular physiology, early knockdown of ribosomal genes immediately after infection resulted in the elimination of parasites from mice **(Fig 6G-H, Fig S17A)**, likely due to impaired stem cell function that prevents successful tissue establishment. In contrast, RNAi-mediated interference starting from 14 dpi markedly inhibited parasite growth without reducing worm burden in permissive mouse host **(Fig 6H-J, Fig S17B)**. Collectively, these findings indicate that ribosomal gene down-regulation contributes to translational repression, thereby impeding the progression of proliferative cells into differentiated progeny. This repression likely contributes to the developmental arrest of *S. japonicum* from schistosomula to adult worms in rats.

Loss of ribosomal components and dysfunction in ribosomal activity directly led to impaired peptide synthesis and formation of protein aggregates^[64]^. The proteotoxicity of ribosome-stalled polypeptides could inducing cellular stress responses^[64, 65]^. Under external stimuli or adverse intracellular conditions, organelle stress responses like UPR, ER stress, and mitochondrial stress often exhibited functional crosstalk^[66, 67]^. Peng et al. reported that NO in rats induced ER and mitochondrial stress in *S. japonicum*, and indicated that misfolded proteins may serve as one of the triggers of ER stress^[28]^. NO not only directly disrupted ER function and activated ER stress-mediated apoptotic pathways^[68]^, but also reduced translational activity and caused ribosome collision^[69]^, affecting ribosomal complex assembly, impairing peptidyl transferase activity and inhibiting cell proliferation^[70, 71]^. By disrupting ribosomal function, NO may be responsible for the reduced protein synthesis capacity of *S. japonicum* in rats. The impaired *de novo* protein synthesis observed in schistosomula from rats in this study may represent a sign of ribosomal stress and potentially played an initiating role in the multi-organelle dysfunction induced by rats.

Taken together, this study reveals that the developmental arrest of *S. japonicum* in the non-permissive rat host, in contrast to the permissive mouse host, is initiated between 12 and 24 dpi. Single-cell transcriptomic profiling of parasites from rats and mice further identified specific cell types and gene expression patterns associated with developmental arrest in the rat host. Our results demonstrate that disrupted energy metabolism and impaired ribosomal function are key factors contributing to this arrested development. These findings offer novel insights into the molecular mechanisms governing host-specific developmental trajectories of schistosomes and highlight potential targets for therapeutic intervention and schistosomiasis control.

## Materials and Methods

### Ethics statement

The manipulation of animals was conducted in strict compliance with the animal care and use guidelines established by Fudan University. All procedures were approved by the Animal Care and Use Committee of Fudan University (Fudan IACUC 201802158S) to ensure the ethical and responsible treatment of the animals.

### *S. japonicum* infection and perfusion

*S. japonicum* cercariae were obtained from *Oncomelania hupensis* snails provided by the Anhui Institute of Schistosomiasis Control. For worm collection or *in vivo* RNA interference experiments, 6-week-old female C57 mice (SiPeiFu Biotechnology Co., Ltd.) and 80-120 g female Sprague-Dawley (SD) rats (Beijing Vital River Laboratory Animal Technology Co., Ltd.) were used. Mice were percutaneously infected via the abdominal skin with approximately 200, 100, or 40 cercariae and perfused at 12, 24, or 42 dpi, respectively. Rats were infected with 1000 cercaria percutaneously. Schistosomula or adult worms were collected from mice or rats by perfusion through the superior vena cava with saline solution. Recovered worms were rinsed with DMEM medium containing 10% horse serum, anesthetized with ethyl 3-aminobenzoate methanesulfonate to dissociate worm pairs, and used for worm counting, length measurement, or fixation.

### Parasite staining and labelling

For carmine staining, male and female *S. japonicum* worms were anesthetized and fixed with AFA fixative (24% formaldehyde, 50% ethanol, 4% acetic acid). Worms were then stained with alum carmine solution as previously described^[63]^. Following staining, the testes of male worms and the ovaries and vitellarium of female worms were observed with Nikon A1 laser scanning confocal microscope under the 594 nm laser channel.

To label proliferative cells, *S. japonicum* worms were collected from mice and rats at 12 and 24 dpi. The worms were immediately transferred DMEM supplemented with 10% horse serum-containing and incubated with 10 μM 5-ethynyl-2’-deoxyuridine (EdU) at 37 °C with 5% CO₂ for 4 h to label proliferating cells. EdU detection was performed as previously described^[72]^. Briefly, fixed worms were subjected to a sequential treatment consisting of bleaching, proteinase K digestion, and re-fixation. The samples were then incubated in the dark for 30 minutes in a detection solution comprising 1% 100 mM CuSO₄, 0.1% azide-conjugated fluorescent dye (545 nm), and 20% 500 mM ascorbic acid. Z-stack confocal images were acquired using a Nikon A1 laser scanning confocal microscope with excitation at 545 nm.

To label nascent proteins, worms collected from mice and rats at 14 dpi, as well as from *in vitro* RNAi and control groups, were immediately transferred into BM169 medium containing 10 μM O-propargyl-puromycin. Worms were incubated at 37°C with 5% CO₂ for 4, 10, 13 or 24 h to label nascent proteins. Following incubation, worms were fixed and processed for visualization using the same protocol employed for EdU detection. During fluorescence intensity analyses, excitation light intensity and imaging parameters were held constant across all samples. Z-stack confocal images were merged using the average intensity projection method in ImageJ. Relative OPP fluorescence intensity, normalized to DAPI staining, was quantified to assess differences in protein synthesis capacity between worms from different hosts or experimental groups.

### Scanning and transmission electron microscope

Fresh *S. japonicum* worms were washed with PBS and fixed in a glutaraldehyde-containing fixative at 4 °C in the dark. Fixed samples were then submitted to Wuhan Servicebio Technology Co., Ltd. for subsequent embedding, sectioning, and imaging using a scanning electron microscope (SEM) (Hitachi SU8100) and a transmission electron microscope (TEM) (Hitachi HT7800).

### Bulk RNA sequencing and analysis

*S. japonicum* worms were recovered from mice and rats at 12, 24, and 42 dpi, anesthetized, washed with PBS, rapidly frozen in liquid nitrogen, and stored at -80 °C. Samples of three biological replicates were proceeded by Novogene Co., Ltd. for total RNA extraction, transcriptome library preparation and RNA sequencing on Illumina platform.

Gene expression levels (FPKM) were used for PCA with the Subread software, and *k*-means clustering analysis of gene expression was conducted using the pheatmap package in RStudio. Differential expression analysis was performed using DESeq2 (v1.20.0), and significantly up- or down-regulated genes (*p* ≤ 0.05, | Log_2_FoldChange | ≥ 1) were selected to conduct Gene Ontology (GO) and KEGG enrichment analysis using clusterProfiler (v3.8.1). Gene Set Enrichment Analysis (GSEA) was performed in RStudio. Genes selected with *p* ≤ 0.05 and | Log_2_FoldChange | ≥ 0.5 were further analyzed for cell type enrichment using the LRcell package in RStudio, with gene enrichment score matrix derived from previous scRNA-seq data of adult *S. japonicum*.

### Single-cell RNA sequencing and analysis

Male *S. japonicum* worms were perfused from mice and rats at 14 and 24 dpi, washed with PBS, and transferred into 37 °C cell preservation solution. Subsequent single cell dissociation and 10× Genomics single-cell RNA sequencing was conducted by Genergy Bio-Technology Co., Ltd. Two biological replicates were included for each host at each time point. Raw sequencing data were aligned to the *S. japonicum* reference genome assembly version 3 (PRJNA739049) using CellRanger software. Seurat objects were constructed in RStudio using Seurat and hdf5r packages. Quality control retained cells with 200-5000 detected genes and mitochondrial gene content below 15%. After merging samples and removing batch effects, UMAP dimensionality reduction was performed on the integrated Seurat object.

Based on the single-cell atlas of *S. mansoni*^[32]^, marker genes for 23 cell types of *S. japonicum* were identified **(Table S2)**, and 17 cell types were annotated from UMAP clusters. Overlaid UMAP plots were generated for samples from rats and mice at 14 and 24 dpi. Cell density heatmaps and ridge plots were created using the MASS and fields packages, respectively. Cell proportion data were extracted for qualitative and quantitative comparisons of proportion differences between worms from mice and rats.

Gene expression levels for each cell cluster of worms from rats compared to worms from mice at 14 and 24 dpi were calculated with negbinom method, generating lists of DEGs. Relative expression levels of essential DEGs in each cell type was visualized using DotPlot function as bubble plots.

Reference marker genes for the S phase and G2/M phase of the cell cycle in *S. japonicum* were identified through homologous gene screening **(Table S9)**. In RStudio, Seurat subsets of major proliferative cell types (somatic stem cells, tegument progenitors and progeny 1/2, and muscle progenitors) of 14 dpi worm samples from mice and rats were used for cell cycle score calculation with CellCycleScoring function in Rstudio. S phase and G2/M phase scores were stored in the metadata, and the relative proportion of each cell cycle phase was calculated for quantitative comparison.

### *in vitro* treatment of uncoupling agents

*S. japonicum* worms recovered from mice at 14 and 24 dpi were transferred into 12-well plates containing 3 mL BM169 medium and cultured *in vitro* at 37 °C with 5% CO₂. After a 24 h acclimation period, 3 μM oligomycin or 4.5 μM FCCP was added to the treatment groups, while 0.3% anhydrous ethanol (drug solvent) was added to the control group. Worms were continuously cultured for 48 h, after which images were captured and samples were collected and frozen for further analysis.

### Quantitative real-time PCR

Total RNA from worms was extracted using AG RNAex Pro Reagent (Accurate Biotechnology (Hunan) Co., Ltd), following the manufacturer’s instructions. cDNA was synthesized using the Hifair® III 1st Strand cDNA Synthesis SuperMix for qPCR (gDNA digester plus) (Yeasen Biotech Co., Ltd., Shanghai, China). qPCR was performed using the LightCycler 96 Real-Time PCR System (Roche) following the standard two-step protocol, with *S. japonicum 26S proteasome non-ATPase regulatory subunit 4* (*Sjpsmd*) used as the internal reference. Relative expression levels of target genes were calculated using the 2^-ΔΔCt^ method. Primer sequences used for qPCR are listed in **Table S10**.

### Protein sequences alignment

The amino acid sequence of *Sj*L-LDH and its homologs L-LDH_A in model organisms including *Caenorhabditis elegans*, *Drosophila melanogaster*, *Danio rerio*, *Mus musculus*, *Rattus norvegicus*, and *Homo sapiens* were downloaded from the UniProt database for multiple sequence alignment using the MUSCLE algorithm in MEGA v10.2.6. The alignment results were visualized using the online tool ESPript 3.0 with default settings. Corresponding sequences and IDs are listed in **Table S7**.

### Western blot

Total protein was extracted from *S. japonicum* male worms recovered from mice and rats at 14 and 24 dpi using T-PER Tissue Protein Extraction Reagent (Thermo Fisher Scientific), and protein concentration was measured with Bradford method. PAGE gels (12.5%) were prepared using a rapid PAGE gel preparation kit (Yazyme Biopharma, Shanghai). Western blot was performed using 15 μg of protein per sample. LDHA Rabbit Polyclonal Antibody (Beyotime Biotechnology, Shanghai) was used to detect *Sj*L-LDH, and β-Actin was detected using mouse monoclonal β-Actin antibody (Affinity Biosciences). Goat anti-mouse or anti-rabbit secondary antibodies (Affinity Biosciences) were used accordingly. Grayscale intensity of the 35 kD *Sj*L-LDH bands relative to the 42 kD β-Actin bands were quantified in ImageJ to compare the relative expression levels of *Sj*L-LDH in worms from different hosts.

### in vitro RNAi

Ribosomal gene sequences were cloned into the pJC53.2 plasmid, and double-stranded RNA (dsRNA) was synthesized using the T7 High Yield RNA Synthesis Kit (Yeasen Biotechnology, Shanghai), following the manufacturer’s protocol. Control dsRNA was generated from a PCR-amplified fragment of the pJC53.2 plasmid using T7 primers. *S. japonicum* schistosomula were isolated from infected mice at 14 dpi and cultured *in vitro*, with approximately 30 worms per well in 3 mL of BM169 medium in 12-well plates. Following a 24-hour acclimatization period, the culture medium was replaced every two days, and dsRNA was added at a final concentration of 30 μg/mL. Each RNA interference and control condition was conducted in triplicate. After 22 days of *in vitro* treatment, worms were imaged and collected for further analysis. Primers used for amplification of target regions are listed in **Table S10**.

### in vivo RNAi

Beginning on the day of *S. japonicum* cercarial infection (0 dpi), mice in both experimental and control groups received intravenous injections of 30 μg dsRNA via the tail vein every four days. At 24 dpi, adult worms were collected by perfusion, and worm burden and length were recorded; samples were subsequently preserved for further analysis. The livers and spleens of the mice were photographed for morphological assessment. For ribosomal gene targeting, an additional experimental group was included in which dsRNA administration commenced at 14 dpi to assess stage-specific effects. Each interference and control group included a minimum of three biological replicates. Primers used for amplification of the interference fragments are listed in **Table S10**.

### Statistical analysis and graphics

Statistical data were quantitatively analyzed with the mean and standard deviation (sd) from the results of multiple biological replicate experiments. Statistical comparisons of experimental data were performed using independent-sample *t*-tests. Plots and *t*-test results were generated in GraphPad Prism. Figures of bulk and scRNA-seq analyses were partially created using RStudio or the online platform Bioinformatics (https://www.bioinformatics.com.cn)^[73]^. Images acquired from stereomicroscopy or confocal fluorescence microscopy were processed using ImageJ. Cell 3D reconstruction and quantification were conducted with Imaris v9.0.1.

## Supporting Information

**Fig S1. *k*-means clustering of gene expression across developmental stages.** Heatmap showing *k*-means clustering of gene expression profiles in transcriptomes of *S. japonicum* samples collected from mice and rats at different developmental stages. Abbreviations: C, worms from C57 mice; S, worms from SD rats; M, male worms; F, female worms.

**Fig S2. Expression patterns of DEGs in *S. japonicum* recovered from rats compared with those from mice at 12 and 24 dpi across distinct cell types.**

**Fig S3. UMAP clustering and cell type annotation of worm samples from mice and rats at 14 and 24 dpi. A**, All 60 cell clusters on UMAP plot of integrated Seurat object. **B**, Cell type annotation of 60 cell clusters. Abbreviations: ES, esophageal glands.

**Fig S4. UMAP visualization and cell density distribution across samples. A-D**, UMAP heatmaps showing single-cell transcriptomic profiles of *S. japonicum* at 14 dpi from mice (**A**) and rats (**B**), and at 24 dpi from mice (**C**) and rats (**D**), with notable cell types highlighted by boxed regions. Blue shading indicates relative cell density at specific locations on the UMAP plot. Ridge plots display the distribution of cell types, with colors representing distinct tissue types and ridge height indicating relative cell density at corresponding positions. Abbreviations: *S.j*, *S. japonicum*; C57, worms from C57 mice; SD, worms from SD rats; ES, esophageal glands.

**Fig S5. Cell-level distribution of differentially expressed genes in *S. japonicum* at 14 and 24 dpi from different hosts.** Bar plots showing the number of DEGs at the single-cell level in worms derived from rats compared to those from mice. Upregulated genes in rat-derived worms are shown in red, and downregulated genes in blue. Abbreviations: ES, esophageal glands.

**Fig S6. Cell-level expression changes of mitochondrial DNA-encoded genes in 14 dpi *S. japonicum* from rats compared to those from mice.** Abbreviations: C57, worms from C57 mice; SD, worms from SD rats. ES, esophageal glands.

**Fig S7. Effects of *in vitro* oligomycin and FCCP treatment on mitochondrial gene expression in 14 and 24 dpi worms.** Statistical analyses were performed using independent-sample *t*-tests. ****, *p* ≤ 0.0001; ***, *p* ≤ 0.001; **, *p* ≤ 0.01; *, *p* ≤ 0.05; ns, not significant (*p* > 0.05).

**Fig S8. Expression profile of *SjL-ldh* in the integrated single-cell transcriptomic atlas of *S. japonicum*.** The intensity of blue indicates the relative expression level of *SjL-ldh* across cell types. Abbreviations: ES, esophageal glands.

**Fig S9. Amino acid sequence alignment of *Sj*L-LDH with lactate dehydrogenases from other model organisms.** The species of origin and the position of the first amino acid in each sequence are indicated on the left. Residues with similarity greater than 0.7 across species are highlighted in red, while fully conserved residues are shown with a red background.

**Fig S10. Worm burden in the *in vivo* control and *SjL-ldh* RNAi-treated groups.** Statistical analyses were performed using independent-sample *t*-tests. ****, *p* ≤ 0.0001; ***, *p* ≤ 0.001; **, *p* ≤ 0.01; *, *p* ≤ 0.05; ns, not significant (*p* > 0.05).

**Fig S11. Expression changes of mitochondrial genes in the *in vivo SjL-ldh* RNAi-treated *S. japonicum*.** Statistical analyses were performed using independent-sample *t*-tests. ****, *p* ≤ 0.0001; ***, *p* ≤ 0.001; **, *p* ≤ 0.01; *, *p* ≤ 0.05; ns, not significant (*p* > 0.05).

**Fig S12. Gene Set Enrichment Analysis (GSEA) of *S. japonicum* from rats compared with those from mice at 12 and 24 dpi.** For each comparison group, the top 10 downregulated pathways were selected based on significantly enriched gene sets with a normalized enrichment score (NES) < 0 and an adjusted *p*-value < 0.25. Abbreviations: *S.j*, *S. japonicum*; C57, worms from C57 mice; SD, worms from SD rats.

**Fig S13. Expression changes of ribosome-associated genes in the *in vivo SjL-ldh* RNAi-treated *S. japonicum*.** Statistical analyses were performed using independent-sample *t*-tests. ****, *p* ≤ 0.0001; ***, *p* ≤ 0.001; **, *p* ≤ 0.01; *, *p* ≤ 0.05; ns, not significant (*p* > 0.05).

**Fig S14. Comparison of proliferating cells in *S. japonicum* form mice and rats at 12 and 24 dpi. A**, EdU labeling of proliferating cells in 12 dpi schistosomula and 24 dpi male worms from mice and rats. Scale bars, 20 μm. Representative images from 4 worms. **B**, Quantitative comparison of EdU⁺ cell proportions in 12 dpi schistosomula from two hosts. **C**, Quantitative comparison of EdU⁺ cell proportions in 24 dpi male worms from two hosts. Statistical analyses were performed using independent-sample *t*-tests. ****, *p* ≤ 0.0001; ***, *p* ≤ 0.001; **, *p* ≤ 0.01; *, *p* ≤ 0.05; ns, not significant (*p* > 0.05).

**Fig S15. Nascent protein synthesis in 14 dpi schistosomula from mice and rats.** Worms were incubated *in vitro* with OPP for 4, 10, 13, and 24 hours following recovery from the respective hosts to assess protein synthesis activity. Scale bars, 100 μm.

**Fig S16. Brightfield imaging of 14 dpi *S. japonicum* after *in vitro* control and ribosomal gene RNAi-treatment. Scale bars, 2000 μm.**

**Fig S17. Morphology of *S. japonicum* collected from control and ribosomal gene *in vivo* RNAi-treated groups. A**, Representative images of worms from control and ribosomal gene RNAi groups with the first injection administered at 0 dpi. Scale bars, 2000 μm. **B**, Representative images of worms from control and ribosomal gene RNAi groups with the first injection administered at 14 dpi. Scale bars, 2000 μm.

**Table S1. Differentially expressed genes in *S. japonicum* worms at 12, 24 and 42 dpi from rats compared to worms from mice.**

**Table S2. Marker genes for each cell type identified in the integrated single-cell RNA sequencing (scRNA-seq) dataset of *S. japonicum*.**

**Table S3. Differentially expressed genes in individual cell types and whole worms of 14 dpi male *S. japonicum* schistosomula from mice compared to those from rats based on scRNA-seq analysis.**

**Table S4. Differentially expressed genes in individual cell types and whole worms of 24 dpi male *S. japonicum* schistosomula from mice compared to those from rats based on scRNA-seq analysis.**

**Table S5. Significantly downregulated genes (*p* ≤ 0.05, Log_2_FoldChange ≤ -0.5) identified from both bulk and single-cell RNA sequencing.**

**Table S6. Glucose metabolism-related genes across scRNA-seq samples of 14 dpi and 24 dpi male *S. japonicum* worms.**

**Table S7. Sequence information of *Sj*L-LDH and homologous lactate dehydrogenases from other model organisms used in the phylogenetic analysis.**

**Table S8. Gene Set Enrichment Analysis (GSEA) results of *S. japonicum* from rats compared with those from mice at 12 and 24 dpi.**

**Table S9. Cell cycle marker genes of *S. japonicum* used for cell cycle phase prediction in single-cell RNA sequencing analysis.**

**Table S10. Gene sequences and primers used in this study.**

## Acknowledgments

This research was funded by the National Key Research and Development Program of China (Grant Nos. 2021YFC2300800 and 2021YFC2300803), Fund of Fudan University and Cao’ejiang Basic Research (No.24FCA04). We thank the staff at the National Institute of Parasitic Disease, Chinese Center for Disease Control and Prevention (NIPD, China CDC) for providing the cercaria of *S. japonicum*. We thank BioRender for providing the scientific illustration platform.

## Author contributions

J.P.W., Y.P.W. and W.H. conducted this project. J.P.W. secured funding and supervised the overall research. Y.P.W. performed the bioinformatics analysis and all experiments. E.L.T. provided assistance in electron microscope images and WB results analysis. S.Y.C. and X.C. provided assistance in bioinformatics analysis. W.J.C. provided experimental assistance in nascent protein staining. Y.H. provided experimental assistance in parasite infection and perfusion. Y.P.W. and J.P.W. drafted the manuscript. All authors critically reviewed and approved the final manuscript.

## Data Availability

Data generated in this study are available upon request from the corresponding author. The raw sequencing data reported in this paper have been deposited in NCBI Sequence Read Archive (SRA) database under the accession number PRJNA1264034 for bulk-RNAseq data and PRJNA1264518 for single-cell RNA-sequencing data.

