## Supplementary Figures for "Metabolic and ribosomal dysregulation drive developmental arrest of *Schistosoma japonicum* in non-permissive rats"

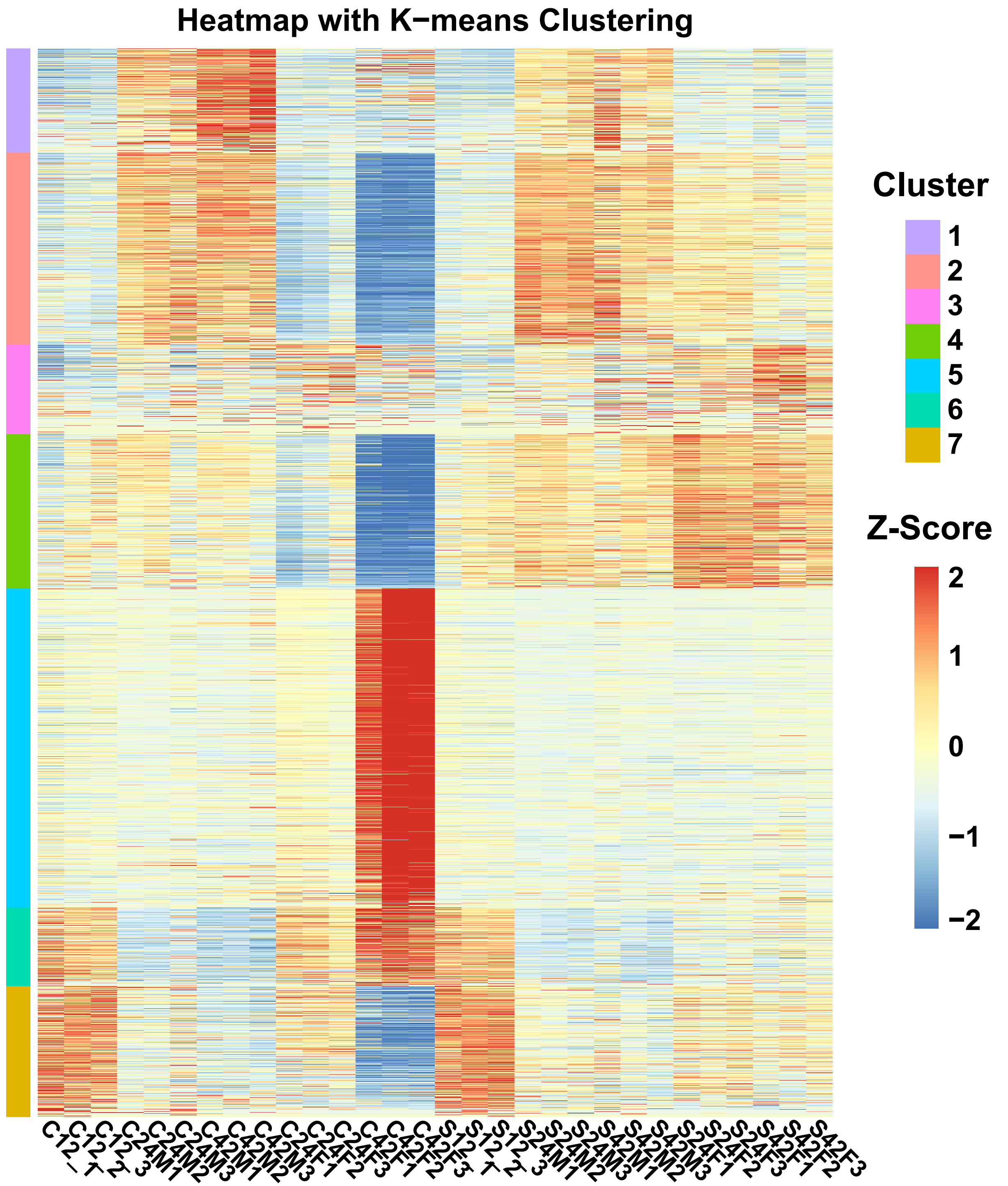


**Supplementary Figure 1. *k*-means clustering of gene expression across developmental stages.** Heatmap showing *k*-means clustering of gene expression profiles in transcriptomes of *S. japonicum* samples collected from mice and rats at different developmental stages. Abbreviations: C, worms from C57 mice; S, worms from SD rats; M, male worms; F, female worms.

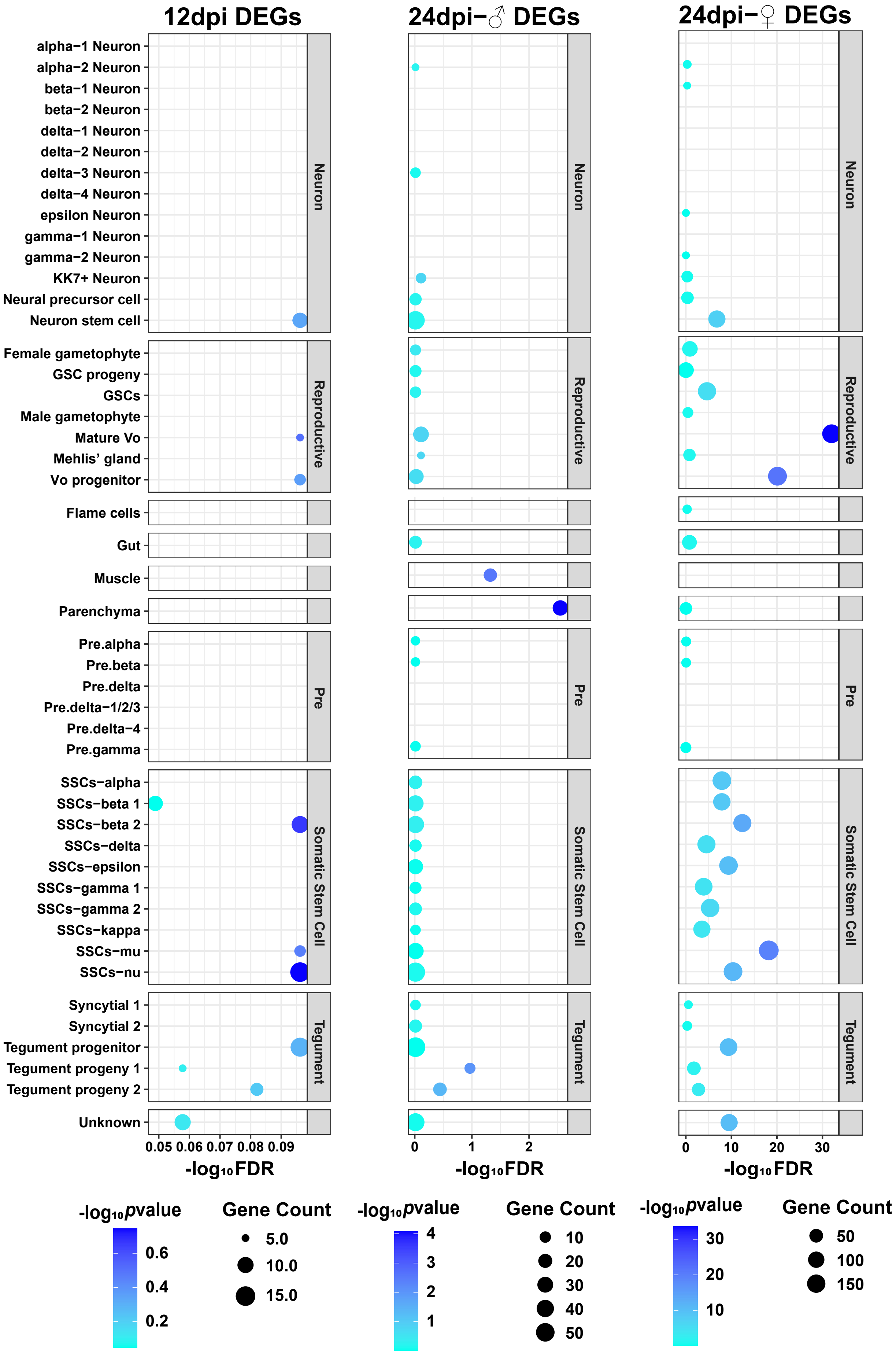


**Supplementary Figure 2.** Expression patterns of DEGs in *S. japonicum* recovered from rats compared with those from mice at 12 and 24 dpi across distinct cell types.

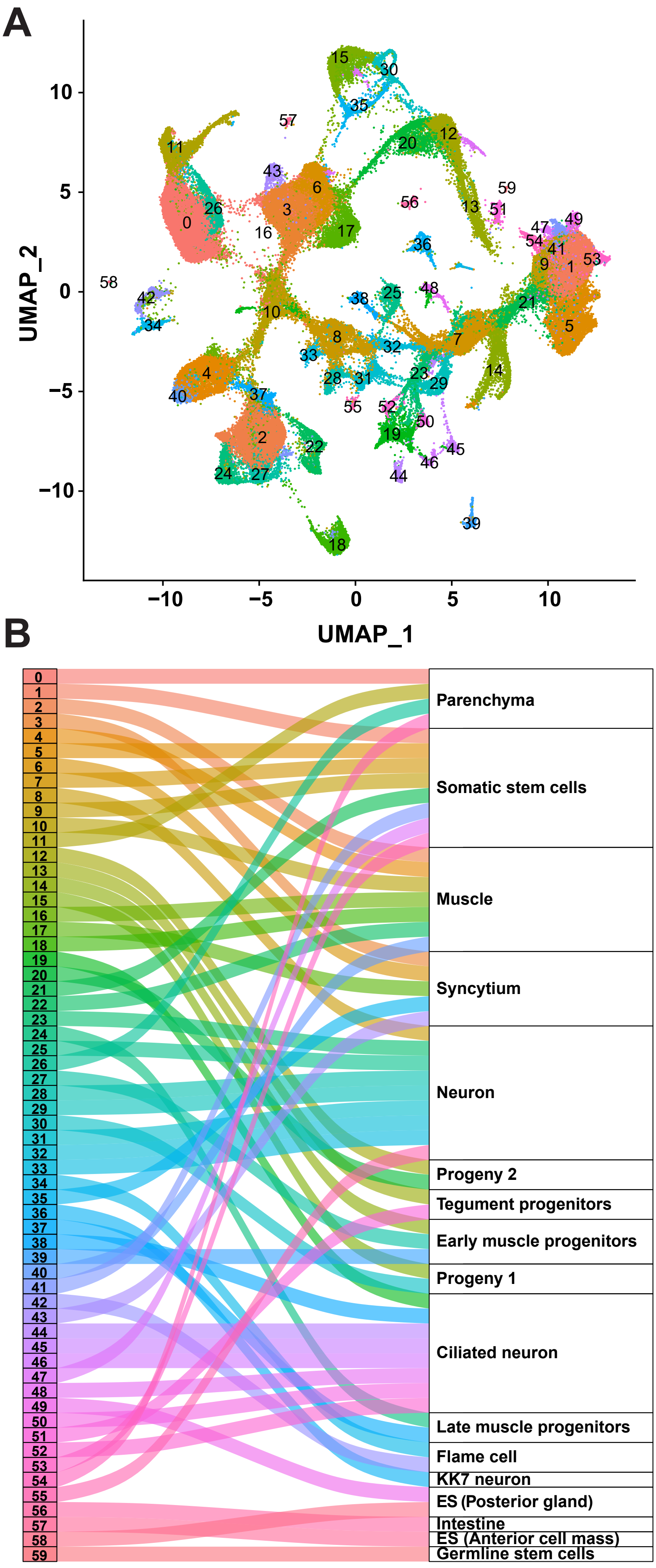


**Supplementary Figure 3. UMAP clustering and cell type annotation of worm samples from mice and rats at 14 and 24 dpi. A**, All 60 cell clusters on UMAP plot of integrated Seurat object. **B**, Cell type annotation of 60 cell clusters. Abbreviations: ES, esophageal glands.

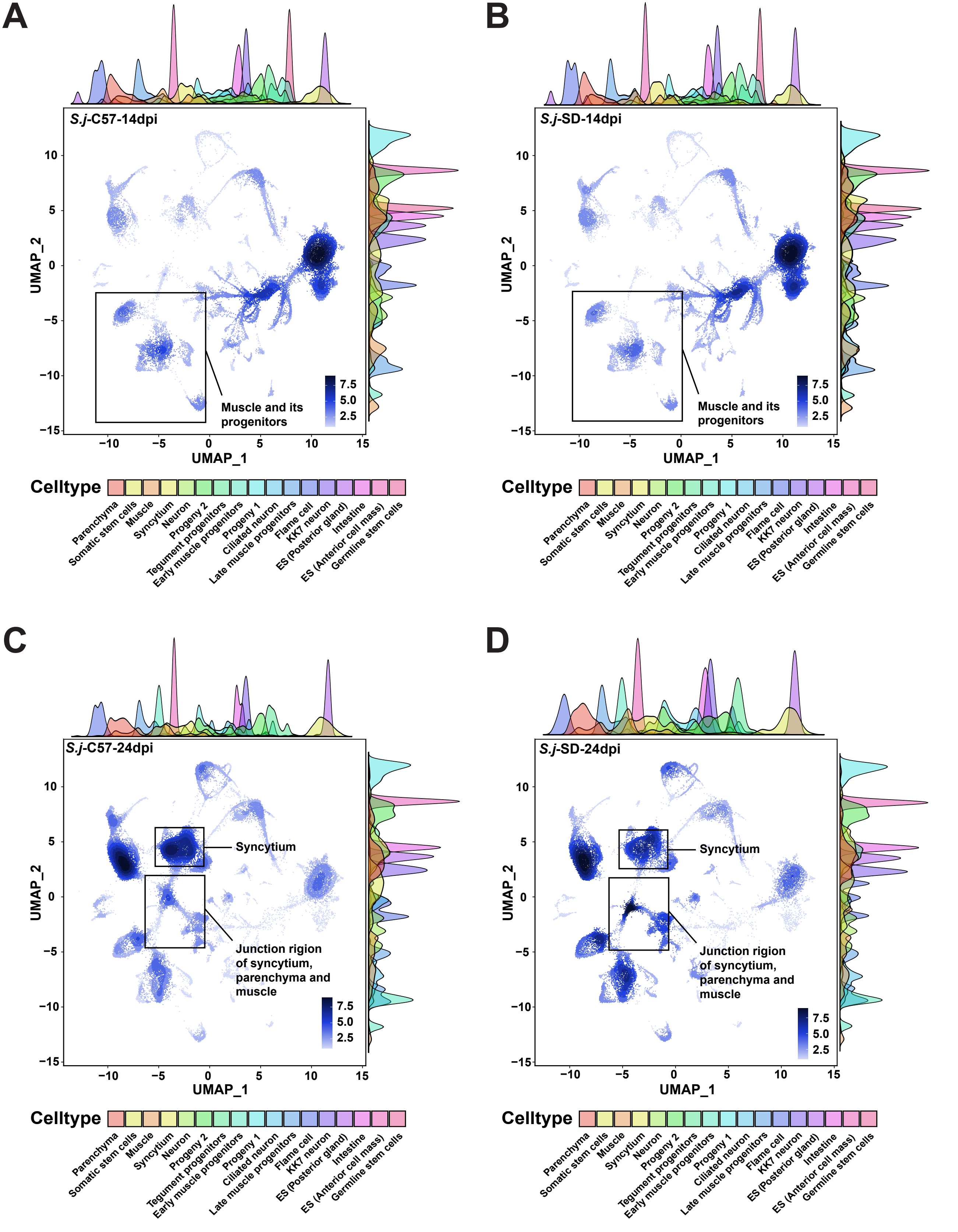


**Supplementary Figure 4. UMAP visualization and cell density distribution across samples. A-D**, UMAP heatmaps showing single-cell transcriptomic profiles of *S. japonicum* at 14 dpi from mice **(A)** and rats **(B)**, and at 24 dpi from mice **(C)** and rats **(D)**, with notable cell types highlighted by boxed regions. Blue shading indicates relative cell density at specific locations on the UMAP plot. Ridge plots display the distribution of cell types, with colors representing distinct tissue types and ridge height indicating relative cell density at corresponding positions. Abbreviations: *S.j*, *S. japonicum*; C57, worms from C57 mice; SD, worms from SD rats; ES, esophageal glands.

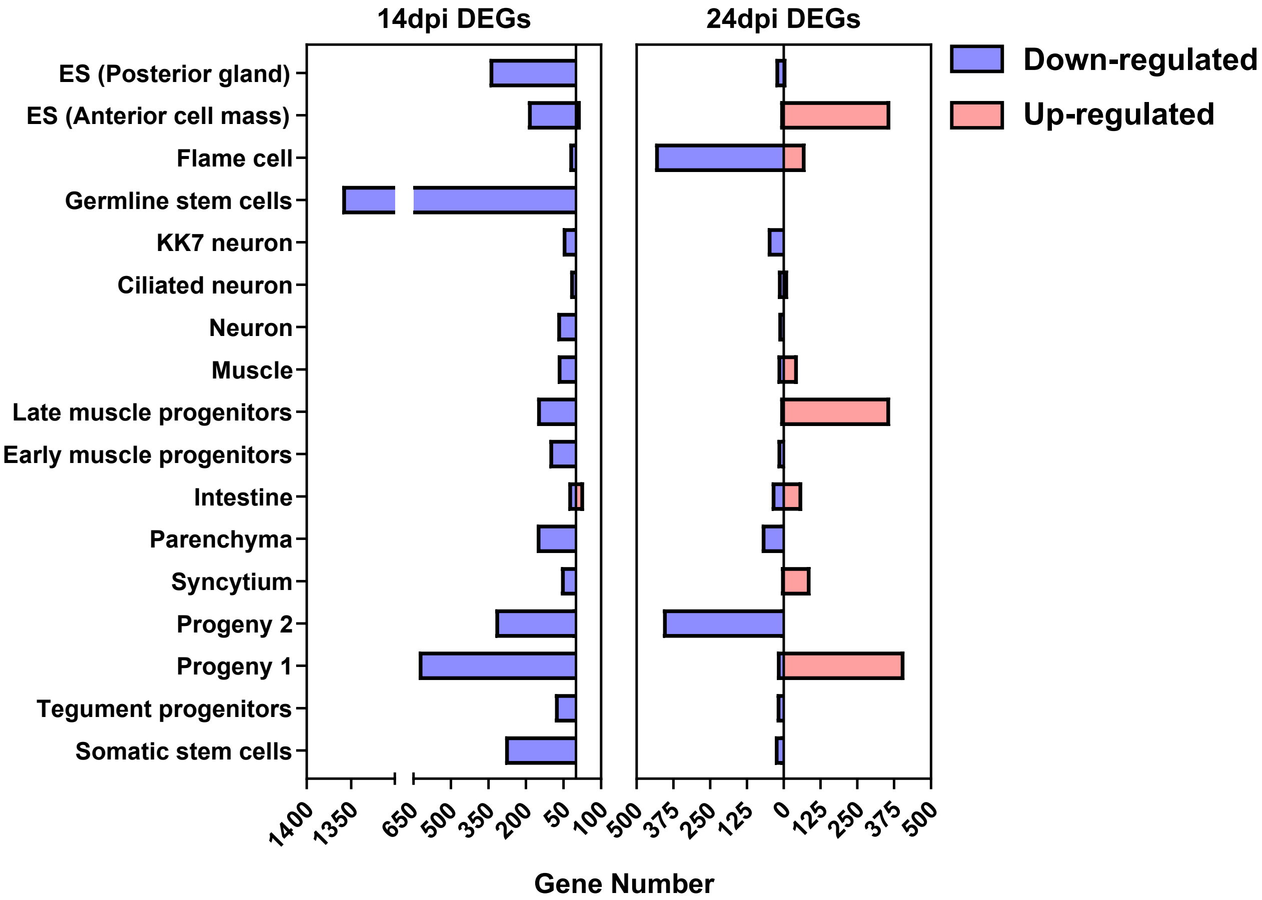


**Supplementary Figure 5. Cell-level distribution of differentially expressed genes in *S. japonicum* at 14 and 24 dpi from different hosts.** Bar plots showing the number of DEGs at the single-cell level in worms derived from rats compared to those from mice. Upregulated genes in rat-derived worms are shown in red, and downregulated genes in blue. Abbreviations: ES, esophageal glands.

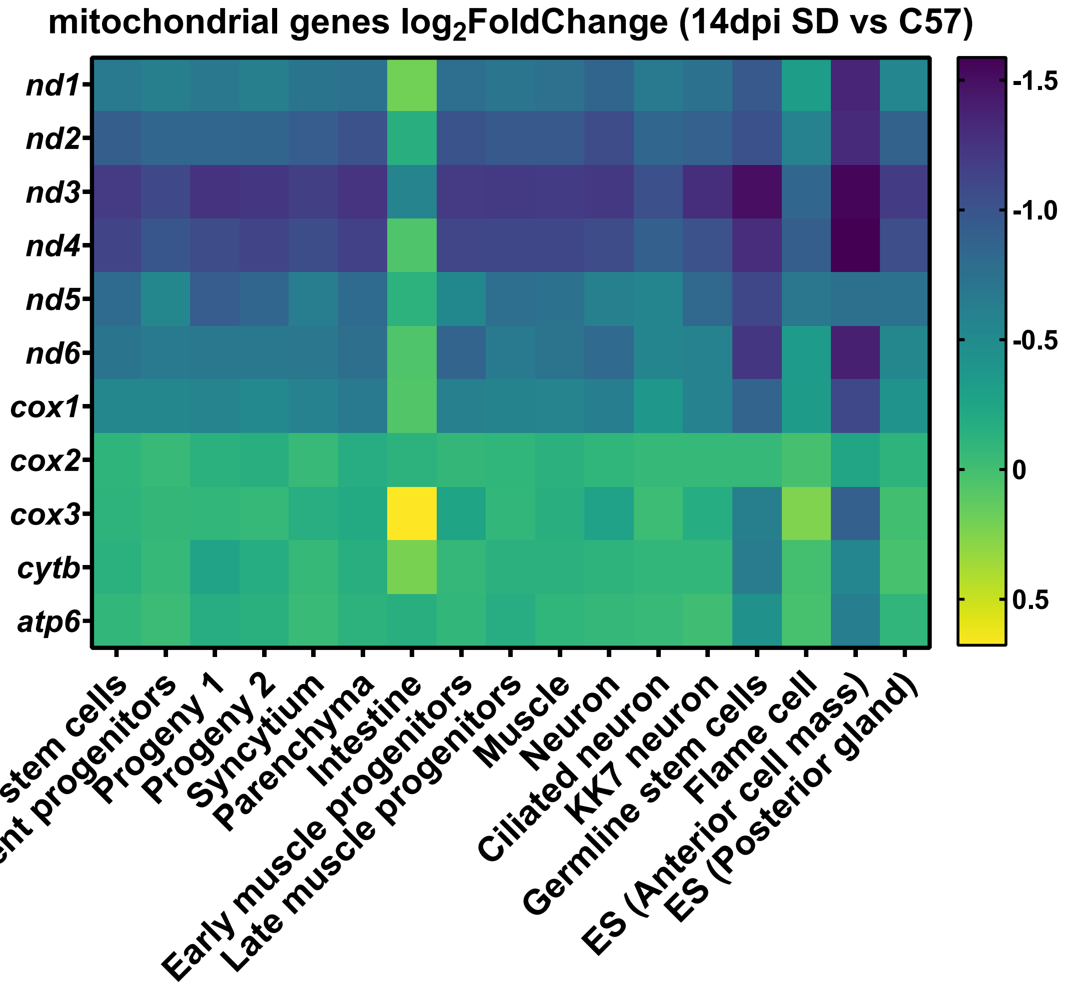


**Supplementary Figure 6. Cell-level expression changes of mitochondrial DNA-encoded genes in 14 dpi *S. japonicum* from rats compared to those from mice.** Abbreviations: C57, worms from C57 mice; SD, worms from SD rats. ES, esophageal glands.

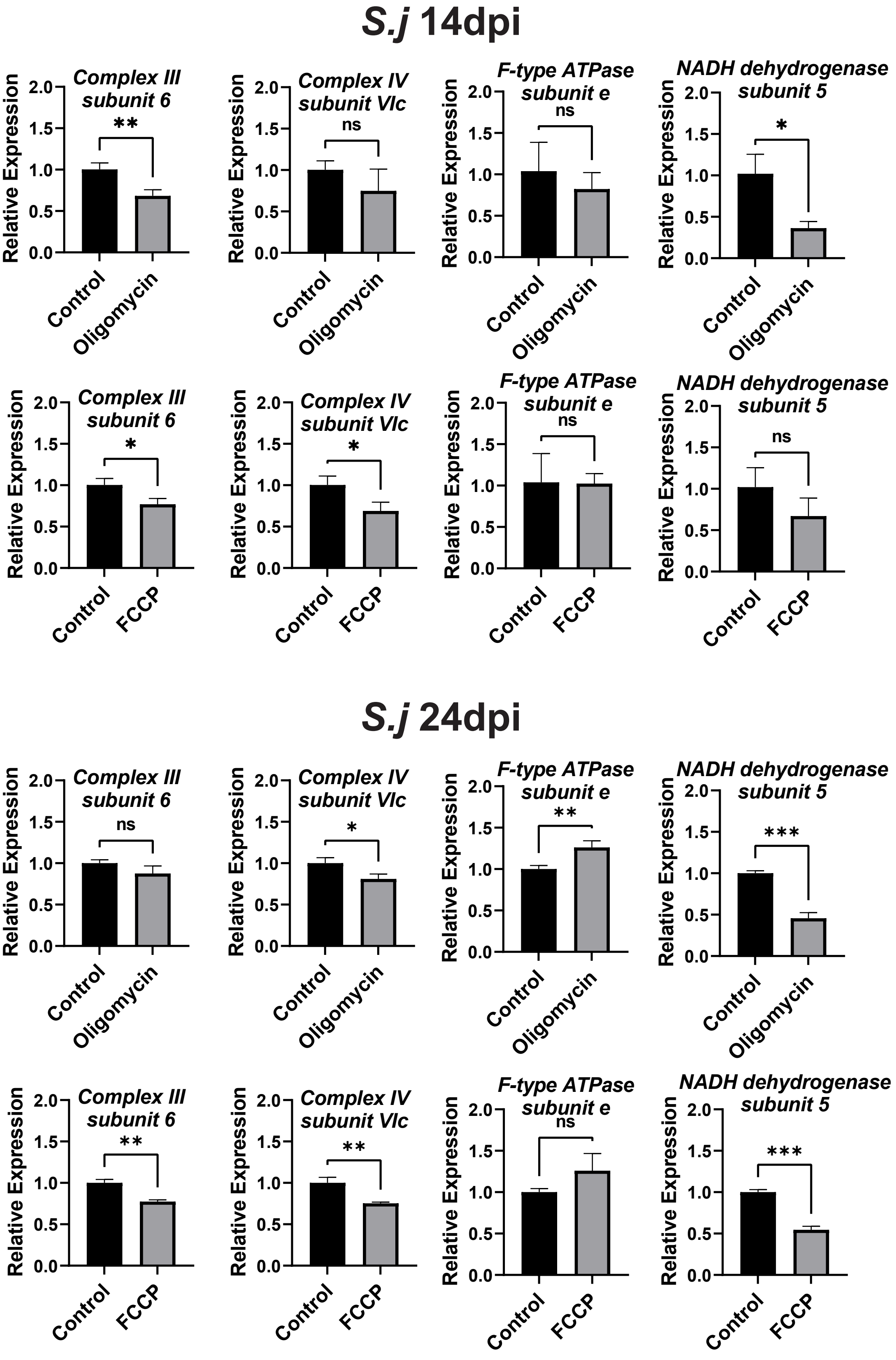


**Supplementary Figure 7. Effects of *in vitro* oligomycin and FCCP treatment on mitochondrial gene expression in 14 and 24 dpi worms.** Statistical analyses were performed using independent-sample *t*-tests. ****, $p \leq 0.0001$; ***, $p \leq 0.001$; **, $p \leq 0.01$; *, $p \leq 0.05$; ns, not significant ($p > 0.05$).

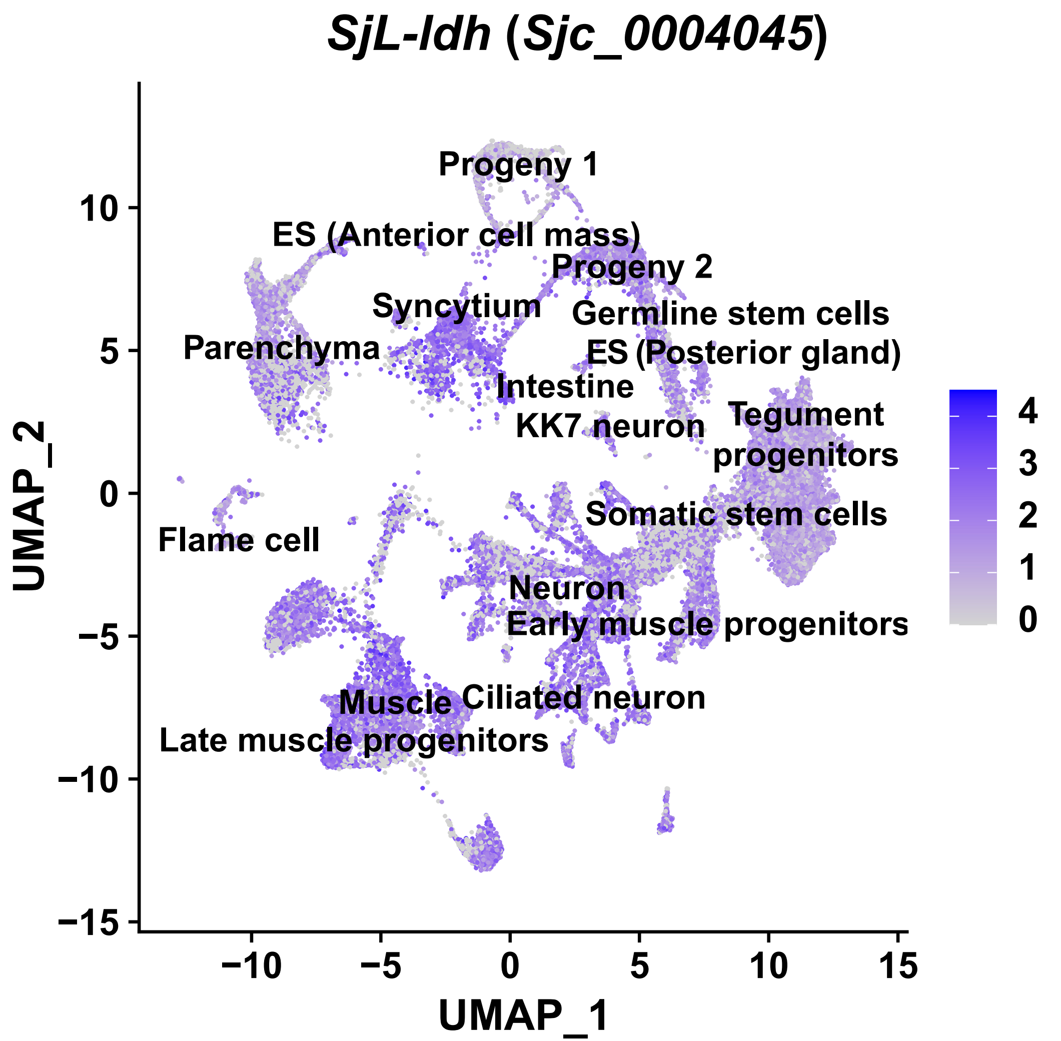


**Supplementary Figure 8. Expression profile of *SjL-ldh* in the integrated single-cell transcriptomic atlas of *S. japonicum*.** The intensity of blue indicates the relative expression level of *SjL-ld*h across cell types. Abbreviations: ES, esophageal glands.

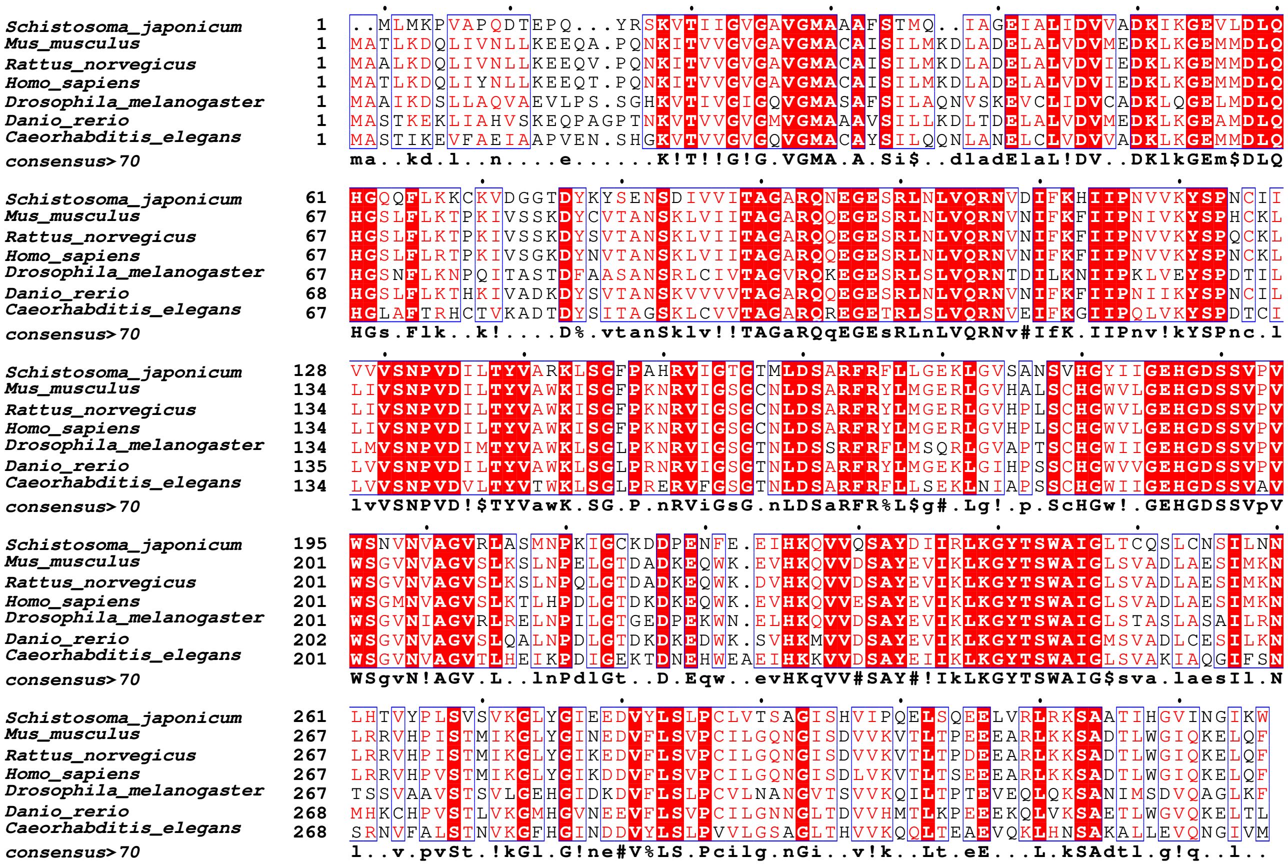


**Supplementary Figure 9. Amino acid sequence alignment of *Sj*L-LDH with lactate dehydrogenases from other model organisms.** The species of origin and the position of the first amino acid in each sequence are indicated on the left. Residues with similarity greater than 0.7 across species are highlighted in red, while fully conserved residues are shown with a red background.

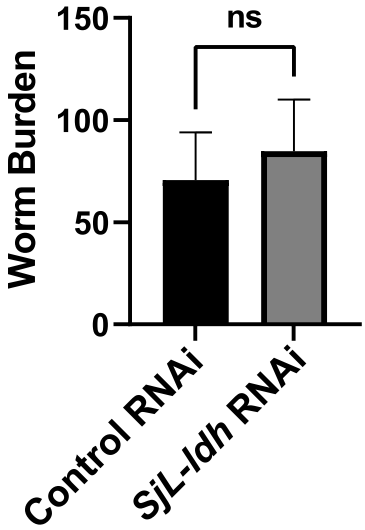


**Supplementary Figure 10. Worm burden in the *in vivo* control and *SjL-ldh* RNAi-treated groups.** Statistical analyses were performed using independent-sample *t*-tests. ****, $p \leq 0.0001$; ***, $p \leq 0.001$; **, $p \leq 0.01$; *, $p \leq 0.05$; ns, not significant ($p > 0.05$).

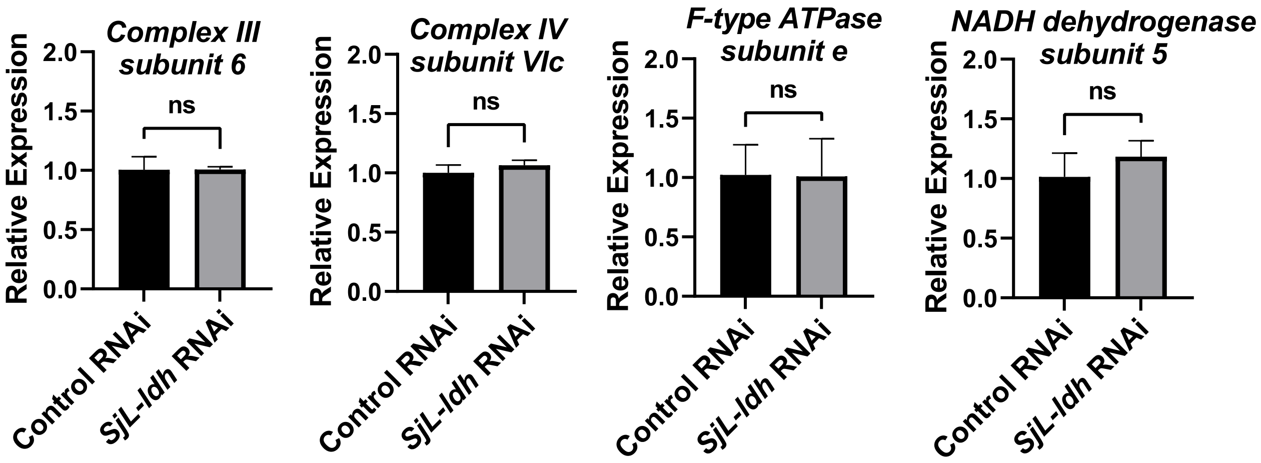


**Supplementary Figure 11. Expression changes of mitochondrial genes in the *in vivo SjL-ldh* RNAi-treated *S. japonicum*.** Statistical analyses were performed using independent-sample *t*-tests. ****, $p \leq 0.0001$; ***, $p \leq 0.001$; **, $p \leq 0.01$; *, $p \leq 0.05$; ns, not significant ($p > 0.05$).

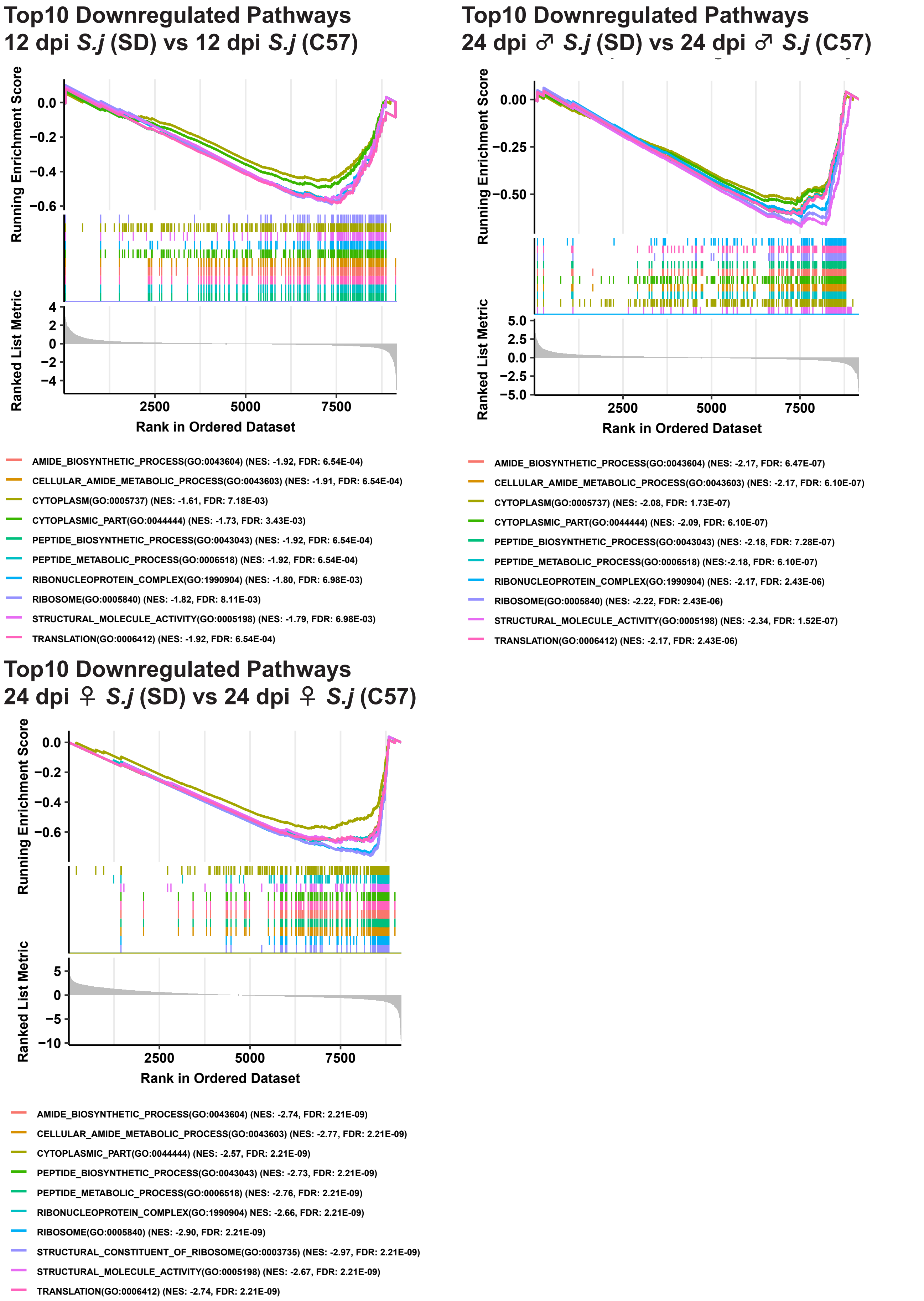


**Supplementary Figure 12. Gene Set Enrichment Analysis (GSEA) of *S. japonicum* from rats compared with those from mice at 12 and 24 dpi.** For each comparison group, the top 10 downregulated pathways were selected based on significantly enriched gene sets with a normalized enrichment score (NES) < 0 and an adjusted *p*-value < 0.25. Abbreviations: *S.j*, *S. japonicum*; C57, worms from C57 mice; SD, worms from SD rats.

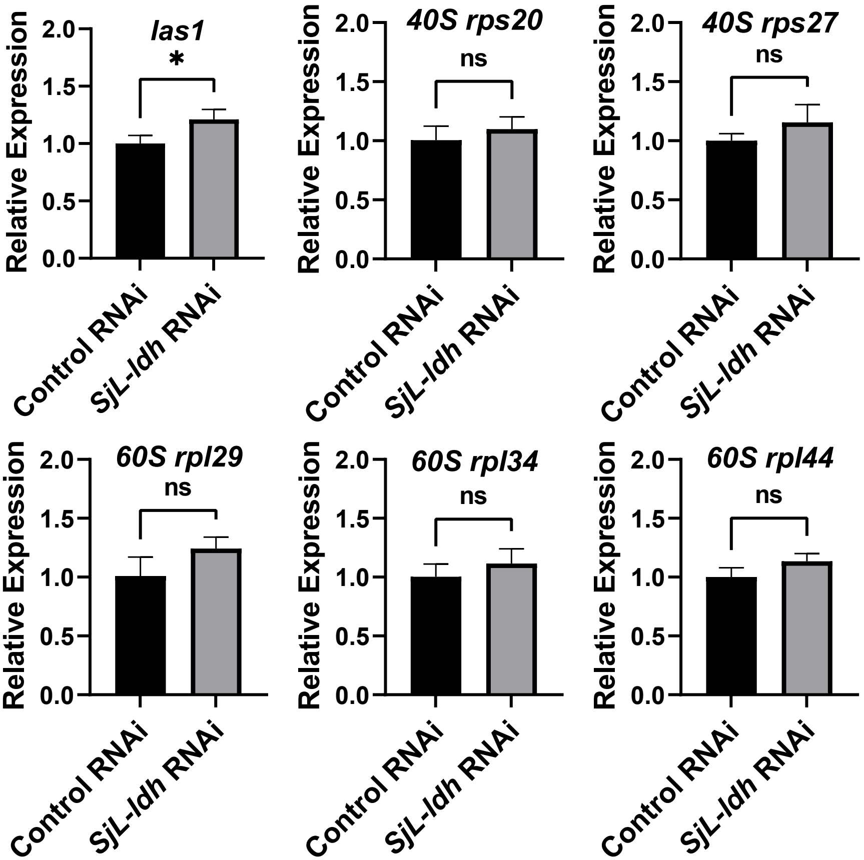


**Supplementary Figure 13. Expression changes of ribosome-associated genes in the *in vivo SjL-ldh* RNAi-treated *S. japonicum*.** Statistical analyses were performed using independent-sample *t*-tests. ****, $p \leq 0.0001$; ***, $p \leq 0.001$; **, $p \leq 0.01$; *, $p \leq 0.05$; ns, not significant ($p > 0.05$).

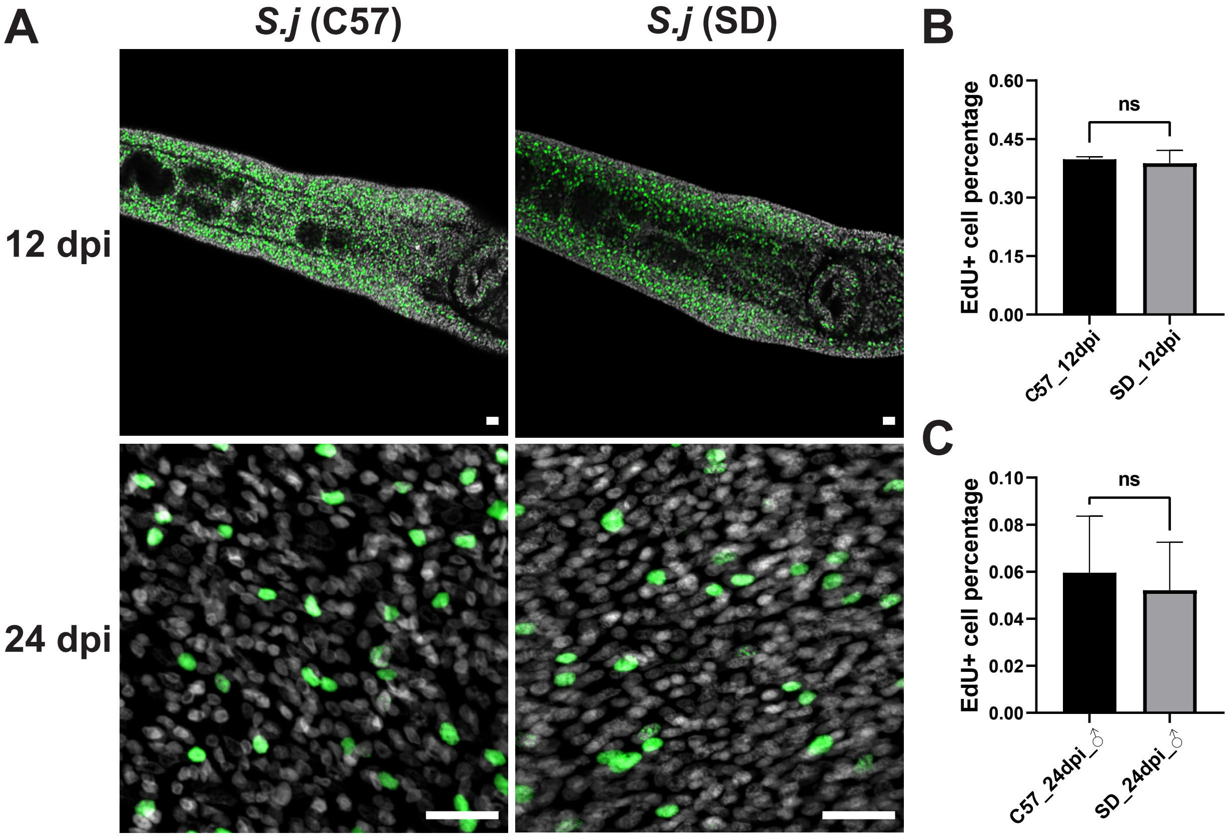


**Supplementary Figure 14. Comparison of proliferating cells in *S. japonicum* form mice and rats at 12 and 24 dpi. A**, EdU labeling of proliferating cells in 12 dpi schistosomula and 24 dpi male worms from mice and rats. Scale bars, 20 µm. Representative images from 4 worms. **B**, Quantitative comparison of $EdU^+$ cell proportions in 12 dpi schistosomula from two hosts. **C**, Quantitative comparison of $EdU^+$ cell proportions in 24 dpi male worms from two hosts. Statistical analyses were performed using independent-sample *t*-tests. ****, $p \leq 0.0001$; ***, $p \leq 0.001$; **, $p \leq 0.01$; *, $p \leq 0.05$; ns, not significant ($p > 0.05$).

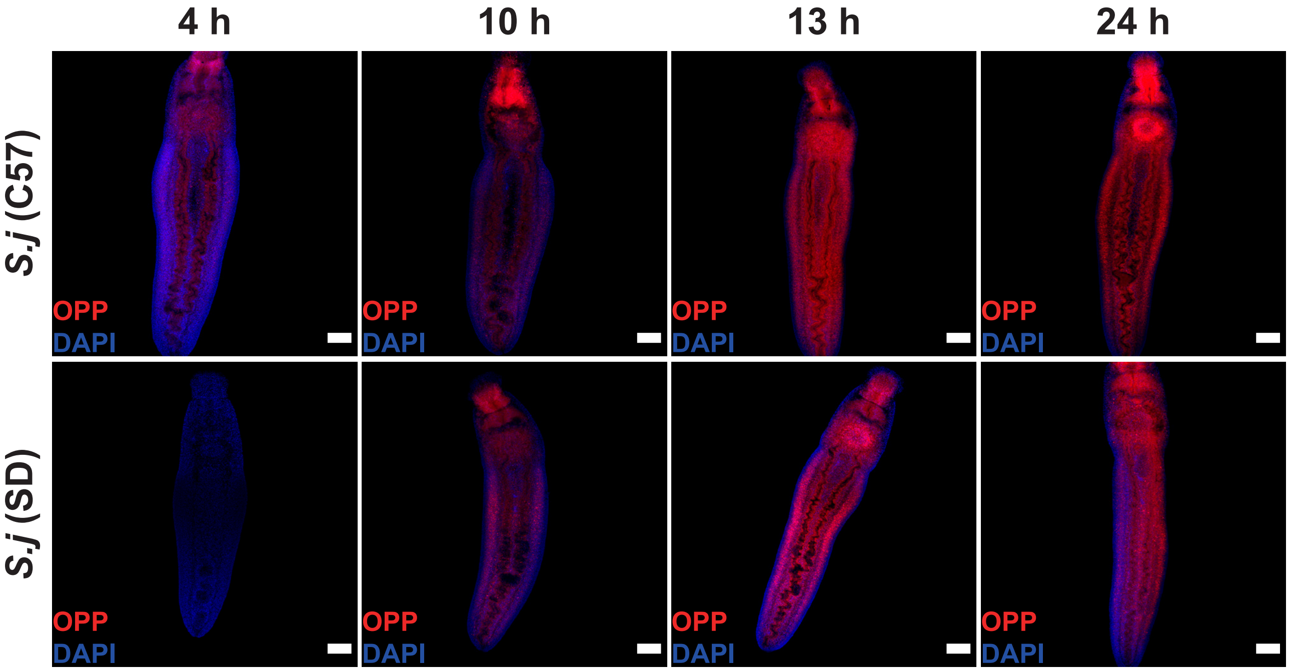


**Supplementary Figure 15. Nascent protein synthesis in 14 dpi schistosomula from mice and rats.** Worms were incubated *in vitro* with OPP for 4, 10, 13, and 24 hours following recovery from the respective hosts to assess protein synthesis activity. Scale bars, 100 μm.

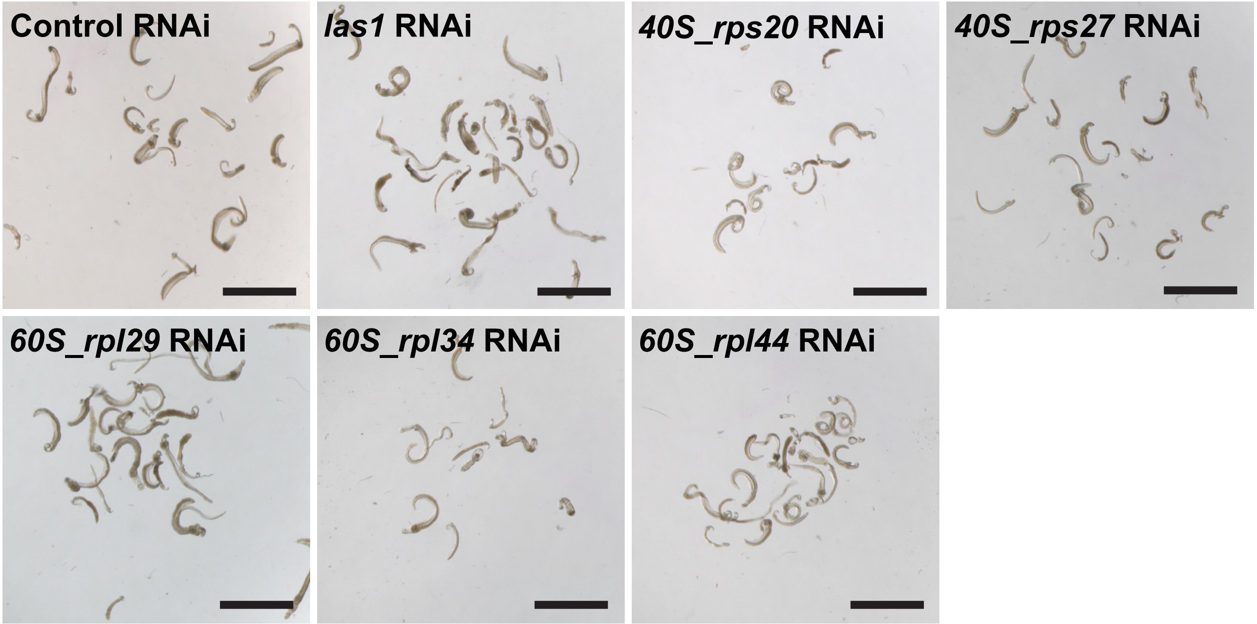


**Supplementary Figure 16. Brightfield imaging of 14 dpi *S. japonicum* after *in vitro* control and ribosomal gene RNAi-treatment.** Scale bars, 2000 μm.

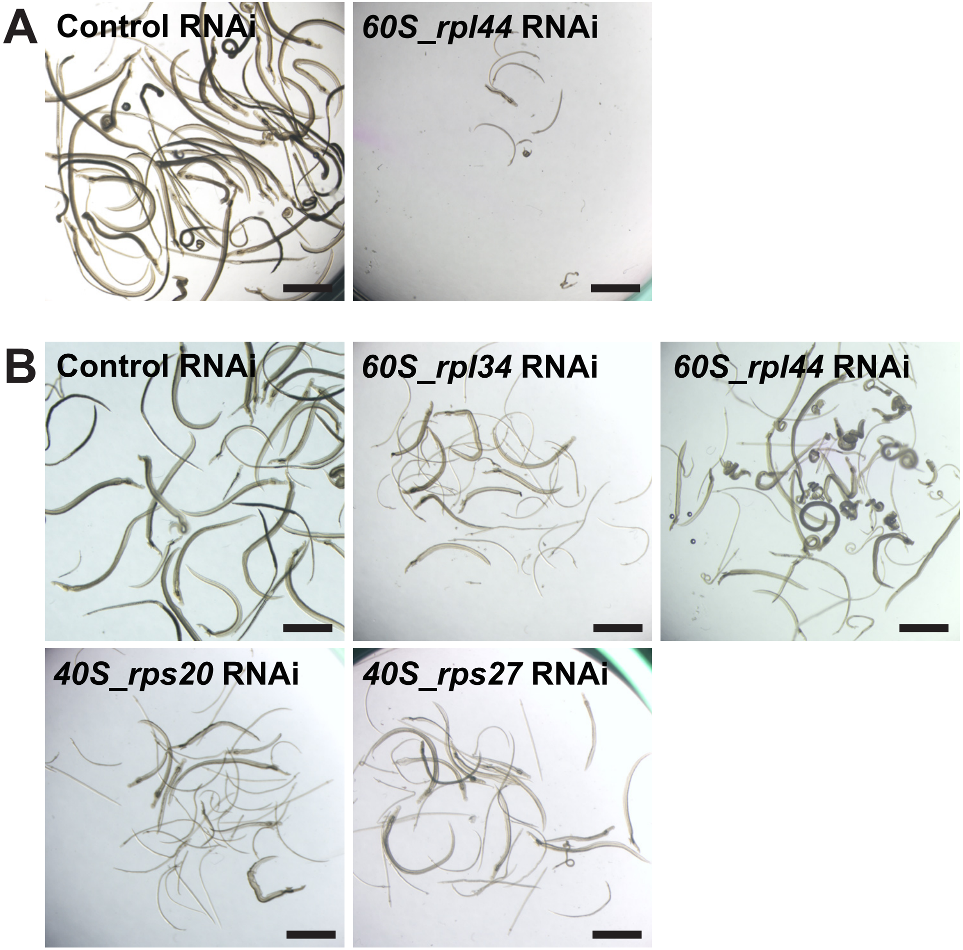


**Supplementary Figure 17. Morphology of *S. japonicum* collected from control and ribosomal gene *in vivo* RNAi-treated groups. A**, Representative images of worms from control and ribosomal gene RNAi groups with the first injection administered at 0 dpi. Scale bars, 2000 μm. **B**, Representative images of worms from control and ribosomal gene RNAi groups with the first injection administered at 14 dpi. Scale bars, 2000 μm.

### Graphic Abstract

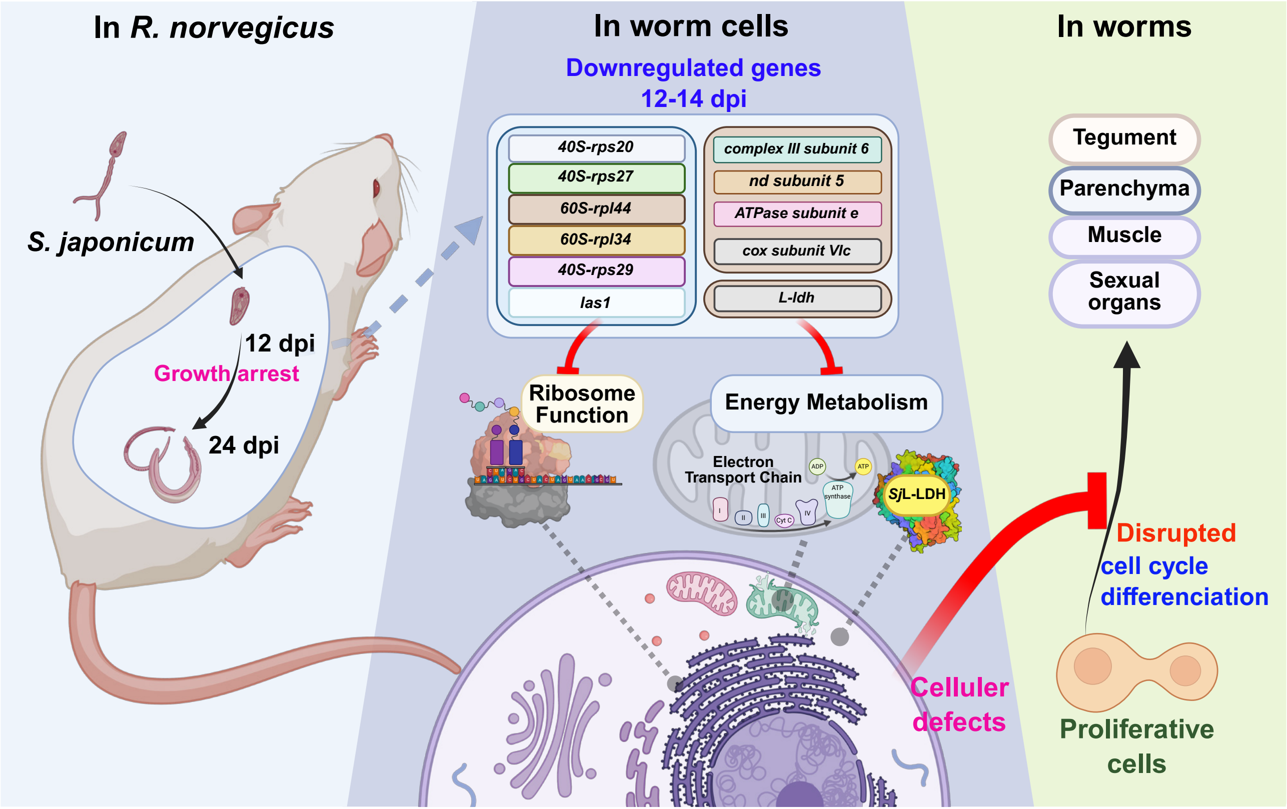
